# Single-nucleus RNA sequencing reveals cellular and molecular dynamics of white and brown adipose tissue in a mouse model of type-2 diabetes

**DOI:** 10.1101/2024.02.06.578860

**Authors:** Taylah L. Gaynor, Ian Hsu, Crisdion Krstevski, Gabriella E. Farrugia, Malathi S. I. Dona, Charles D. Cohen, Brian G. Drew, Alexander R. Pinto

## Abstract

Excessive adipose tissue expansion is often linked with type-2 diabetes. Despite recent efforts mapping adipose tissue changes in obesity using single-cell omics, an understanding of cellular and gene expression changes in a model of type 2 diabetes, and the transcriptional circuitry controlling it, is still lacking. Here, we use single-nucleus RNA sequencing to analyze the remodeling of gonadal white and interscapular brown adipose tissue from female and male mice with or without diabetes. Analysis of 51,877 nuclei revealed altered phenotypes in every cell population in type 2 diabetes. This included an immunoregulatory response, and changes in extracellular matrix, energy generation, and hormone responses. Key transcription factors were inferred as cell-specific and non-specific nodes controlling diabetes-linked phenotypes. Finally, female-to-male population heterogeneity and gene expression differences were observed. Here we provide a resource detailing how adipose tissue remodeling, and the molecular mechanisms governing it, may contribute to cardiometabolic disease.

## Introduction

Cardiometabolic diseases, such as type-2 diabetes mellitus (T2D), have been linked with excessive expansion of adipose tissue^1^. Adipose depots can be broadly classified as white and brown adipose tissue (WAT and BAT, respectively), with WAT being recognized as an endocrine organ having a primary function in lipid storage while BAT drives thermogenesis. The anatomical location of each adipose depot can dictate the tissue’s impact systemically^2^. Specifically, the expansion of visceral WAT can exert pathogenic effects through the release of pro-inflammatory adipokines^3^. Conversely, subcutaneous WAT and interscapular BAT can be protective^4,5^ with research actively focusing on inducing ‘browning’ in WAT to improve energy expenditure and metabolic outcomes^6^.

Many single-cell and spatial transcriptomics studies have provided detailed characterization of adipose depot cellular heterogeneity^7–13^ including how this is altered in obesity^14–21^. Moreover, disease-specific cell populations in obese WAT from individuals with and without T2D have been identified^7,21^. Yet, despite these advancements, a comprehensive analysis of both WAT and BAT in an animal model of T2D, without the influence of confounding variables, is lacking. Further, with biological sex being more widely recognized as a factor influencing disease processes including cardiovascular disease^22^, the role of sex in adipose tissue remodeling remains unclear.

To study the remodeling of WAT and BAT in T2D, we used single-nucleus transcriptomics to profile the entire adipose cellular landscape of female and male mice with or without T2D (*db/db* and *db/h,* respectively). Single-nucleus RNA-sequencing (snRNA-seq) analyses of these depots revealed complex cellular landscapes, identifying important cell-specific gene expression patterns and regulatory circuits that govern cell identity and phenotype. We found that T2D significantly alters the relative abundance of adipose cell types and gene expression of virtually every adipose cell type, resulting in inflammation and tissue remodeling. Further, we identified sex-specific differences in gene expression and cellular abundance, providing a possible basis for varying adipose accretion. Together, these findings provide a valuable resource for understanding WAT and BAT cellular landscapes, their changes in T2D and their potential contribution to metabolic disorders.

## Results

### Leptin-receptor deficient *db/db* mice display hyperglycemia and increased fat mass

To compare adipose depots and characterize adipose remodeling induced by diabetes, we analyzed both female and male *db/h* and leptin-receptor deficient *db/db* mice. To validate hyperglycemia in *db/db* animals, blood glucose levels over an extended time were measured using an HbA1c test (Supplementary Table 1). Higher blood glucose levels were seen in *db/db* mice when compared to *db/h* controls across both females and males (*P*<0.0001 for both) with no sex-specific differences being observed between the genotypes. Next, EchoMRI measurements of lean and fat mass relative to total body weight were taken to assess body composition. EchoMRI showed a greater lean mass in males across both *db/h* and *db/db* genotypes (*P*=0.0036 and *P*=0.0042, respectively). Both sexes had lower lean mass and greater fat mass in *db/db* counterparts (*P*<0.0001 for all). Finally, *db/db* females had greater fat mass than male counterparts (*P*=0.0017). These findings highlight T2D-related hyperglycemia and increased fat mass in *db/db* mice.

### Novel high throughput nuclei isolation technique to characterize the adipose cellulome

Single-cell and spatial transcriptomics has been instrumental in shedding new light on tissue cellularity and disease responses in multiple tissues, including adipose^7–11,16–19,21^. Due to the lipid-filled nature of mature adipocytes, a key approach for single-cell transcriptomic profiling of adipose involves crushing tissue to extract nuclei using a dounce homogenizer^12,13,19^. While this technique has proven useful for nuclei extraction, a limitation of this approach is the number of samples that can be processed promptly. To address this issue, we developed a nuclei isolation technique to isolate both mature adipocyte and stromal vascular fraction (SVF) nuclei from samples in a high-throughput manner. The technique involves the machine-based mechanical dissociation of snap-frozen adipose tissue, and subsequent removal of cellular debris using fluorescence-activated cell sorting. BAT and WAT were homogenized at 30 Hz for 30 and 50 seconds, respectively, in 10-second intervals to isolate the greatest number of nuclei (Supplementary Figure 1A). As a result, this technique produced high-quality, intact nuclei that enabled the identification of most adipose tissue cell populations through snRNA-seq (Figure 1A-C; Supplementary Figure 1A-B).

**Figure 1.**
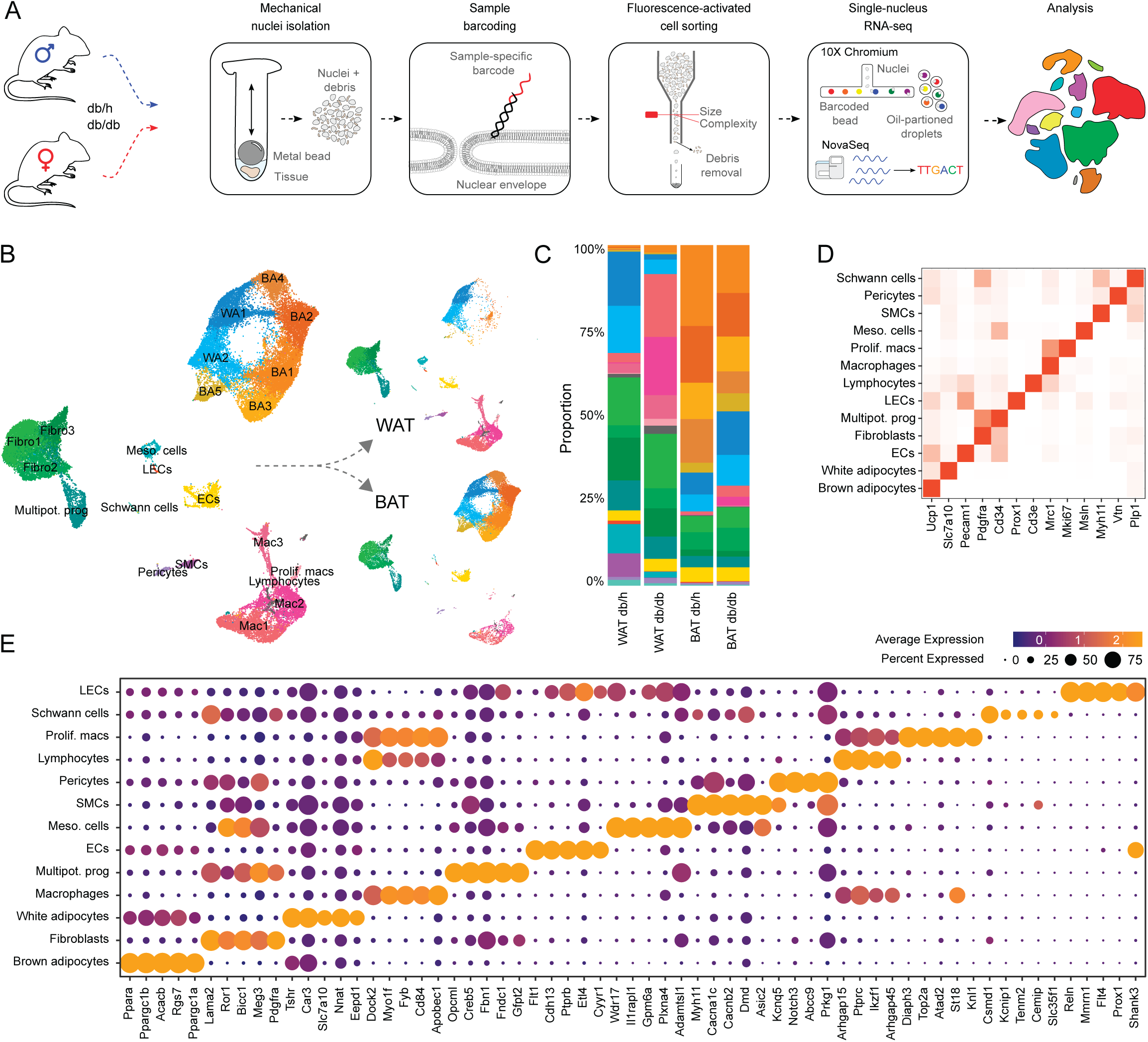
snRNA-seq characterization of adipose cellulome. **(A)** Experimental outline for novel isolation of mature adipocyte and SVF nuclei from snap-frozen adipose tissue. **(B)** UMAP of nuclei representing the adipose cellulome for both *db/h* and *db/db* female and male white and brown adipose tissues. **(C)** Compositional bar chart showing proportion of each cell population in each tissue type. **(D)** Canonical markers of broad adipose tissue cell populations represented in heat map with relative expression indicated by red intensity. **(E)** Top 5 marker genes for each broad cell population with dot size indicating the proportion of cells within that population expressing the marker and dot color indicating the average level of expression. WAT, white adipose tissue; BAT, brown adipose tissue; Fibro, fibroblast; Multipot. prog., multipotent progenitors; Meso. cells, mesothelial cells; ECs, endothelial cells; LECs, lymphatic endothelial cells; SMCs, smooth muscle cells; Mac, macrophage; Prolif. macs, proliferating macrophages; BA, brown adipocyte; WA, white adipocyte.

### snRNA-seq profiling and flow cytometry reveal cellular heterogeneity of adipose depots

Using the aforementioned protocol to isolate nuclei for subsequent snRNA-seq, we were able to generate a dataset containing 51,877 nuclei comprising 22,526 nuclei from gonadal WAT and 29,351 nuclei from interscapular BAT after quality control filtering (Figure 1B). WAT and BAT demonstrated distinct cellular compositions in both *db/h* and *db/db* conditions (Figures 1B-C). Cell populations were identified using their top enriched genes and established marker genes uncovering the cellular heterogeneity of both tissues (Figures 1D-E). Populations identified were macrophages (*Adgre1*, *Fcgr1*, *Csf1r, Cd68*, *Mrc1*), including a proliferating subpopulation (*Top2a*, *Aurka*, *Mki67*), lymphocytes (*Thy1*, *Cd19*, *Cd5*, *Cd3e*), endothelial cells (ECs; *Pecam1*, *Ly6c1*), lymphatic endothelial cells (LECs; *Cldn5, Prox1*), mesothelial cells (*Msln*), smooth muscle cells (SMCs; *Acta2*, *Myh11*), Schwann cells (*Plp1*, *Kcna1*), pericytes (*Pdgfrb*, *Vtn*), multipotent progenitors (*Cd34*, *Dpp4*, *Pi16*, *Creb5*), white adipocytes (*Slc7a10*), and brown adipocytes (*Ucp1*, *Cidea*). Brown adipocytes identified in WAT likely correspond to beige or brite adipocyte subpopulations but are referred to collectively as brown adipocytes throughout analysis. Finally, as fibroblasts and adipocyte progenitors/pre-adipocytes share many of the same marker genes, cells expressing *Pdgfra* and *Col1a1* were collectively referred to as fibroblasts. Epididymal cells (*Adam7*, *Pemt*, *Abcb5*, *Ces5a*), ovarian epithelial cells (*Pax8*, *Epcam*, *Krt7*, *Cdh1*) and skeletal muscle cells (*Ttn*, *Myl1*) were identified and subsequently removed from analysis alongside clusters with high mitochondrial RNA content and low RNA counts (see Methods).

### Adipose cell populations show tissue-dependent compositional and transcriptional individuality

To next compare and validate the cellular compositions of *db/h* WAT and BAT, flow cytometric analysis was conducted on SVF cell populations with mature adipocytes being excluded due to their fragile and buoyant characteristics (Figure 2A; Supplementary Figure 1C). Populations were labelled according to their cell surface marker expression (Supplementary Table 1). Due to challenges in distinguishing adipocyte progenitors, preadipocytes, and fibroblasts from one another, in addition to the lack of clear cell surface markers for these populations^23^, they were collectively labelled as fibroblasts based on their negative expression of mural cell markers. Distinct compositional differences were observed between WAT and BAT. This included variations in most immune and mural cell populations with higher contributions to the viable cell count being observed in WAT when compared to BAT. Of note, vascular EC (VEC) contribution was highest in BAT when compared to WAT, while LECs showed the opposite trend (*P*<0.0001 and *P*=0.0061, respectively). Histological validation and quantification of ECs using isolectin B4 (IB4) showed a greater percentage of IB4 positive (or vascularized) tissue area in BAT corroborating the flow cytometry VEC findings (*P*<0.0001), however, vascularization area per nucleus (stained with DAPI) was not different between depots (Figure 2B).

**Figure 2.**
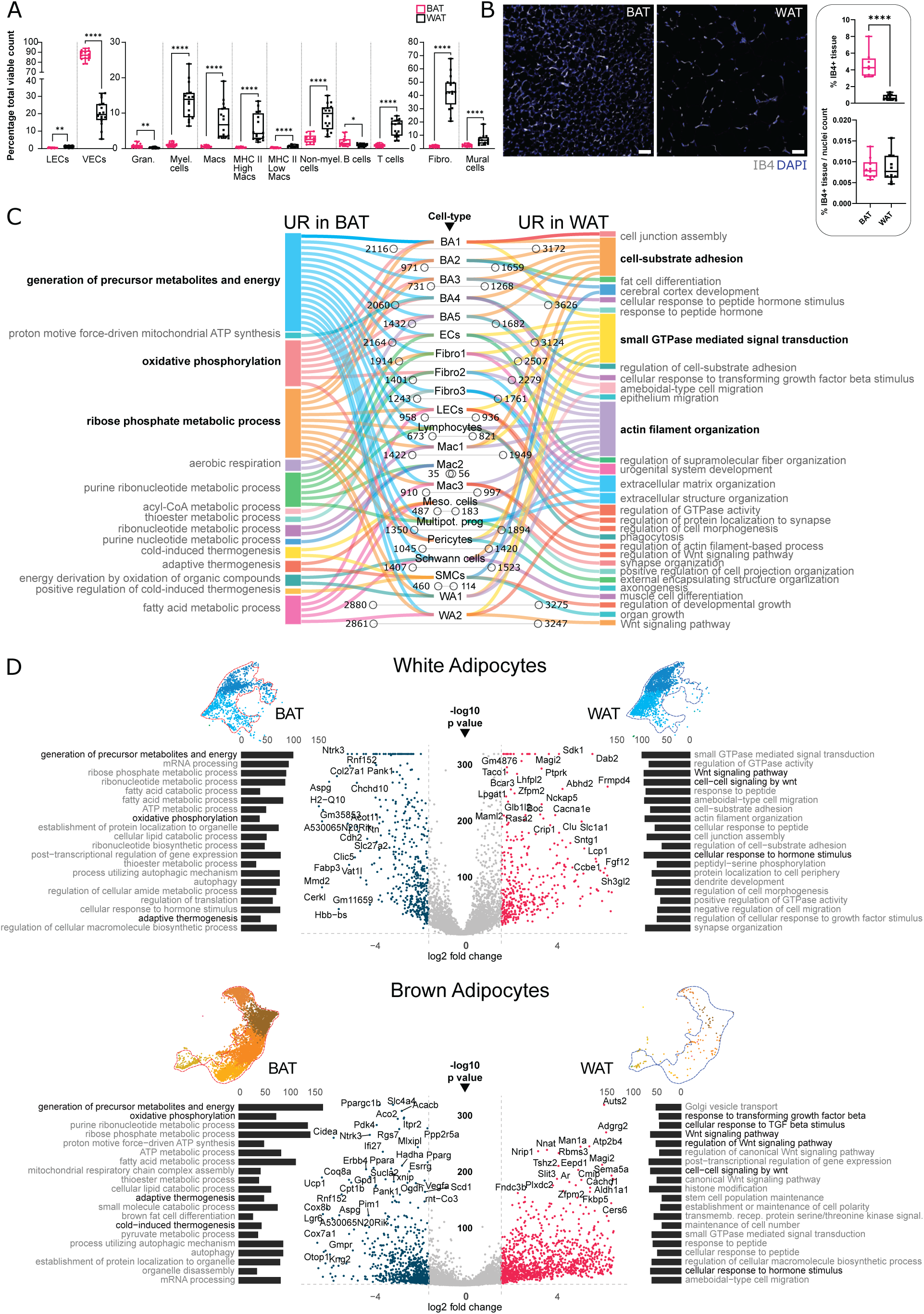
Cellularity and gene expression differences between disparate adipose depots. **(A)** Flow cytometry data of differences in adipose cell population contribution to total viable cell count across BAT (pink) and WAT (black). Statistical significance was assumed at P<0.05. * P< 0.05, ** P< 0.01, *** P< 0.001, **** P< 0.0001. **(B)** Immunofluorescence staining of vasculature (endothelial cells) and nuclei using isolectin B4 (IB4; grey) and DAPI (blue). Scale bar represents 50µm. Statistical significance was assumed at P<0.05. **** P< 0.0001. **(C)** Lollipop plot of upregulated genes across BAT or WAT and Sankey plot of top 3 linked GO terms for each cell population. **(D)** Volcano plots showing differentially expressed genes between white and brown adipocyte subsets in BAT and WAT. Gene placement to the right and left indicate a positive and negative fold difference in expression between tissues, respectively. The placement of the gene on the y-axis indicates significance (P value). GO term bar charts indicating key terms linked to the adjacent genes with the size of the bar representing the number of genes associated.

To assess the gene expression differences between corresponding cell types from WAT and BAT, we performed differential gene expression (DGE) analysis using our snRNA-seq data from *db/h* mice. Our analysis found depot-dependent variations in gene expression across all cell populations with white and brown adipocytes, fibroblasts, and ECs showing the most marked differences (Figure 2C). In BAT, the upregulation of up to 2216 genes in brown adipocytes, 2164 genes in ECs, 1914 genes in fibroblasts, and 2880 genes in white adipocytes was observed and, in WAT, the upregulation of up to 3262 genes in brown adipocytes, 3124 genes in ECs, 2507 genes in fibroblasts, and 3275 genes in white adipocytes was seen. Gene ontology (GO) term analysis of upregulated and downregulated differentially expressed genes (DEGs) uncovered the upregulation of biological pathways in BAT cell populations including *generation of precursor metabolites and energy*, *oxidative phosphorylation*, and *ribose phosphate metabolic process* emphasizing the heat-producing role of this tissue (Figure 2C). GO terms enriched in WAT included *actin filament organization*, *cell substrate adhesion,* and *small GTPase mediated signal transduction.* GTP-mediated RHO signaling has long been implicated in insulin-mediated glucose uptake and homeostasis in adipocytes, highlighting the endocrine role of this tissue^24^. These findings highlight the role of adipose depot-specific microenvironments in programming phenotypes of individual cell populations.

### Adipocyte subpopulation profiles differ between disparate adipose depots

With white and brown adipocytes being present in both WAT and BAT, we subset these nuclei from the dataset and conducted DGE analysis between tissues to determine possible depot-dependent roles (Figure 2D). White adipocytes in BAT showed greater expression of genes including *Fabp3,* of which deficient mice have extreme cold sensitivity^25^. They also showed higher expression of *Ntrk3* (TRKC), a key receptor maintaining cold tolerance and regulating innervation and sympathetic neuron growth in adipose tissue^26^. These genes suggest a possible role for white-like adipocytes in sympathetically activated BAT thermogenic processes. In WAT, white adipocytes were enriched for *Lcp1*, a gene involved in leptin and glucose homeostasis, and in the inhibition of WAT ‘browning’^27^. Expectedly for brown adipocytes in BAT, greater expression of thermogenesis-related genes including *Ucp1*, *Cidea*, *Pparg, Ppara,* and *Ppargc1b* were observed while brown-like adipocytes in WAT were enriched for *Nnat, Aldh1a1*, *Cers6* and *Nrip1* (RIP140), genes, that when suppressed, appear to improve metabolic health via thermogenic pathways^28–31^. GO enrichment analysis of DEGs uncovered pathways such as *generation of precursor metabolites and energy*, *adaptive thermogenesis,* and *oxidative phosphorylation*, among others, as upregulated in adipocytes (irrespective of subtype) from BAT reflecting its heat-producing capacity (Figure 2D). While in WAT, both subpopulations showed expression of genes related to hormonal responses. In addition, terms relating to transforming growth factor beta (TGFβ) and WNT signaling were seen. These data underscore the importance of TGFβ and WNT in driving progenitor proliferation and suppressing adipogenesis^32–34^. WNT signaling GO terms involved genes such as *Tle3* (Supplementary Table 2). TLE3 is a transcription factor co-factor that is selective for white adipocytes by blocking the interaction between PRDM16 and PPARγ and impairing thermogenesis^35^. Together these findings highlight the impact of the cellular landscapes of WAT and BAT on adipocyte phenotypes.

### Distinct gene regulatory circuitry govern brown and white adipose tissue cell phenotypes

Next, we determined the key gene regulatory elements that underlie differences between WAT and BAT and the cells within them. To achieve this, we applied the *single-cell regulatory network inference and clustering* (SCENIC) pipeline^36^. In brief, SCENIC links single-cell sequencing and cis-regulatory data to infer transcriptional modules—regulons—comprising transcription factors (TFs) and their target genes. Cells were first clustered based on their expression of regulons and corresponding gene regulatory networks (GRNs; denoted as TF gene name and ‘(+)’) showing clear clustering patterns that aligned with the previous RNA annotations (Figure 3A; Figure 1B). Moreover, examination of the top 5 regulon markers for each cell population in WAT and BAT demonstrated distinct top GRNs governing cell population phenotypes for brown adipocytes and mesothelial cells, while ECs had shared marker GRNs between tissues with varying levels of similarity observed for other cell populations (Figure 3B). Examples of regulons include Erg(+) and Tal1(+), both important for EC homeostasis and angiogenesis^37,38^, Tfec(+) and Irf5(+), both implicated in macrophage functions and polarisation^39,40^, Meox1(+) and Smad3(+), which govern fibroblast activation^41^, and TGFβ-induced fibrosis^42^. Finally, Esrrg(+), Esrra(+), and Ppara(+) are all implicated in thermogenesis and were observed as markers for brown adipocytes. Altogether these data show an agreement between RNA-seq-based GRN inferences with established transcriptional regulators in various cell populations. This analysis highlights how GRN activity differs between adipose depots to yield tissue-specific cellular phenotypes.

**Figure 3.**
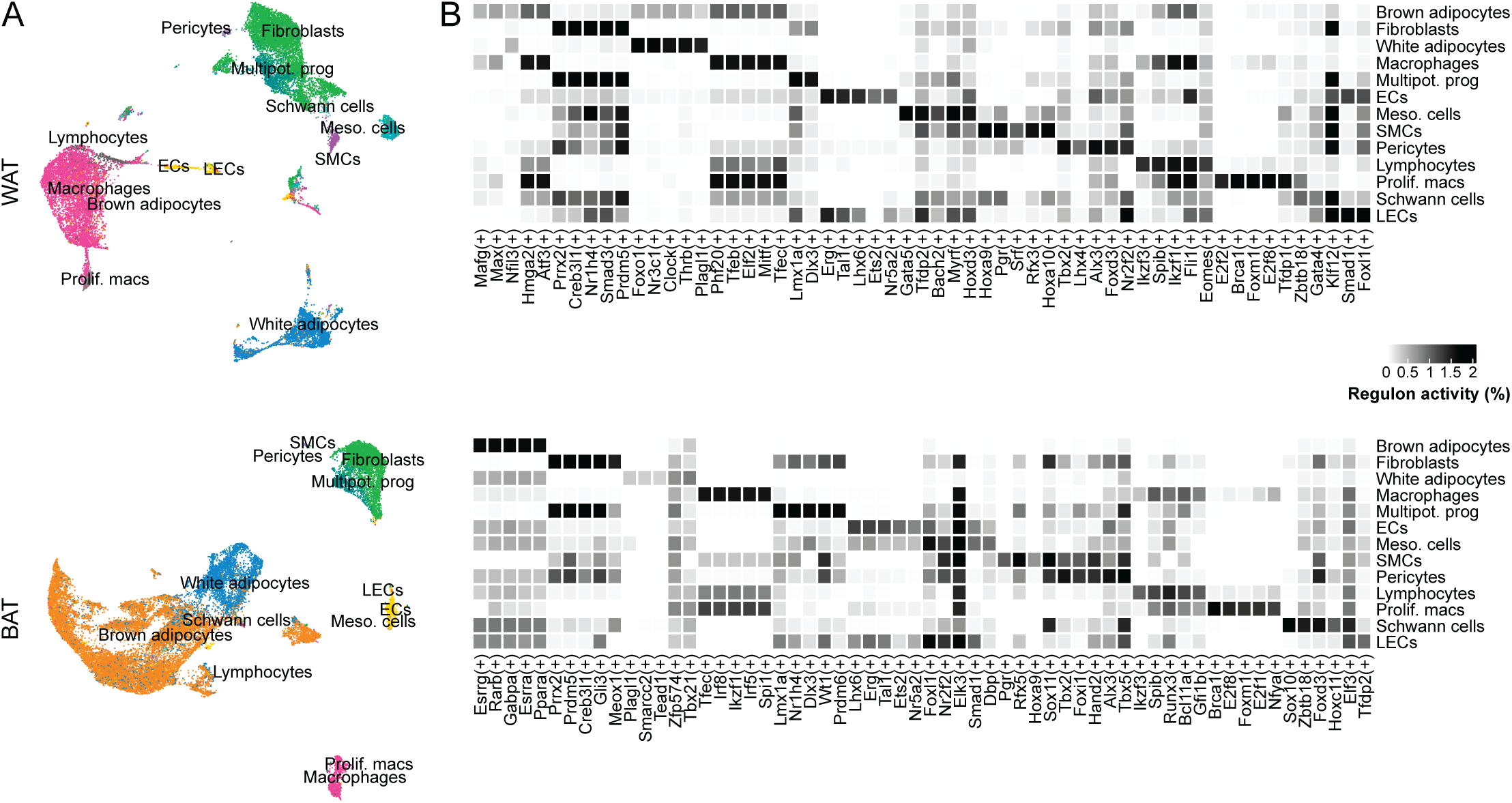
Distinct gene regulatory networks drive cellular phenotypes within brown and white adipose tissue depots. **(A)** UMAP clustering based on regulon-specific expression patterns. Annotation of cell populations carried over from RNA-based clustering seen in Figure 1B. **(B)** Top 5 marker regulons for each cell population based on AUCBinary assay. Intensity of black indicates average regulon activity for each cell population.

### Inflammation and macrophage accumulation in type-2 diabetes adipose depots

To next identify the impact of T2D on WAT and BAT we compared the cellular abundance of *db/h* and *db/db* samples for each depot. Amongst the most striking changes was the increase in macrophages across both WAT and BAT from *db/db* depots (Figure 4A) agreeing with statistical analysis of snRNA-seq counts and flow cytometry data (Supplementary Figure 2). These cellular changes accompanied lipid droplet expansion in adipocytes across both depots of *db/db* mice (Figure 4B). DGE analysis of WAT and BAT between *db/h* and *db/db* tissues identified extensive gene expression differences in virtually all cells, with macrophages amongst those exhibiting the greatest shifts in expression (Figure 4C). In macrophages, the top 3 GO terms associated with upregulated genes for each cell population included *phagocytosis* and *leukocyte migration* (Figure 4C-D). A possible consequence of lipotoxicity and adipocyte hypertrophy is the formation of macrophage crown-like structures (CLSs). CLSs can accumulate in dysfunctional adipose depots through macrophages clustering around dead or dying adipocytes to clear lipid and cell debris^43^. Recent work has highlighted the importance of macrophage proliferation in the formation of these structures and subsequent inflammation^44,45^. Indeed, histological validation using phosphohistone H3 validated the presence of proliferative cells in the CLSs of WAT corroborating the snRNA-seq findings and affirming that macrophage crowns exist in adipose tissue from T2D mice (Figure 4E). Moreover, subpopulation-specific expression of proliferative markers *Top2a*, *Mki67*, and *Aurka* were observed (Figure 4F). Finally, as it is challenging to attribute adipose tissue pathogenesis solely to a T2D-derived phenotype, we assessed the similarity of gene expression patterns independent of obesity between our dataset and a previously published human obesity dataset^21^. Approximately 47% of DEGs across all cell populations in *db/db* mice were unique to our dataset and not differentially expressed in human obesity (Figure 4G).

**Figure 4.**
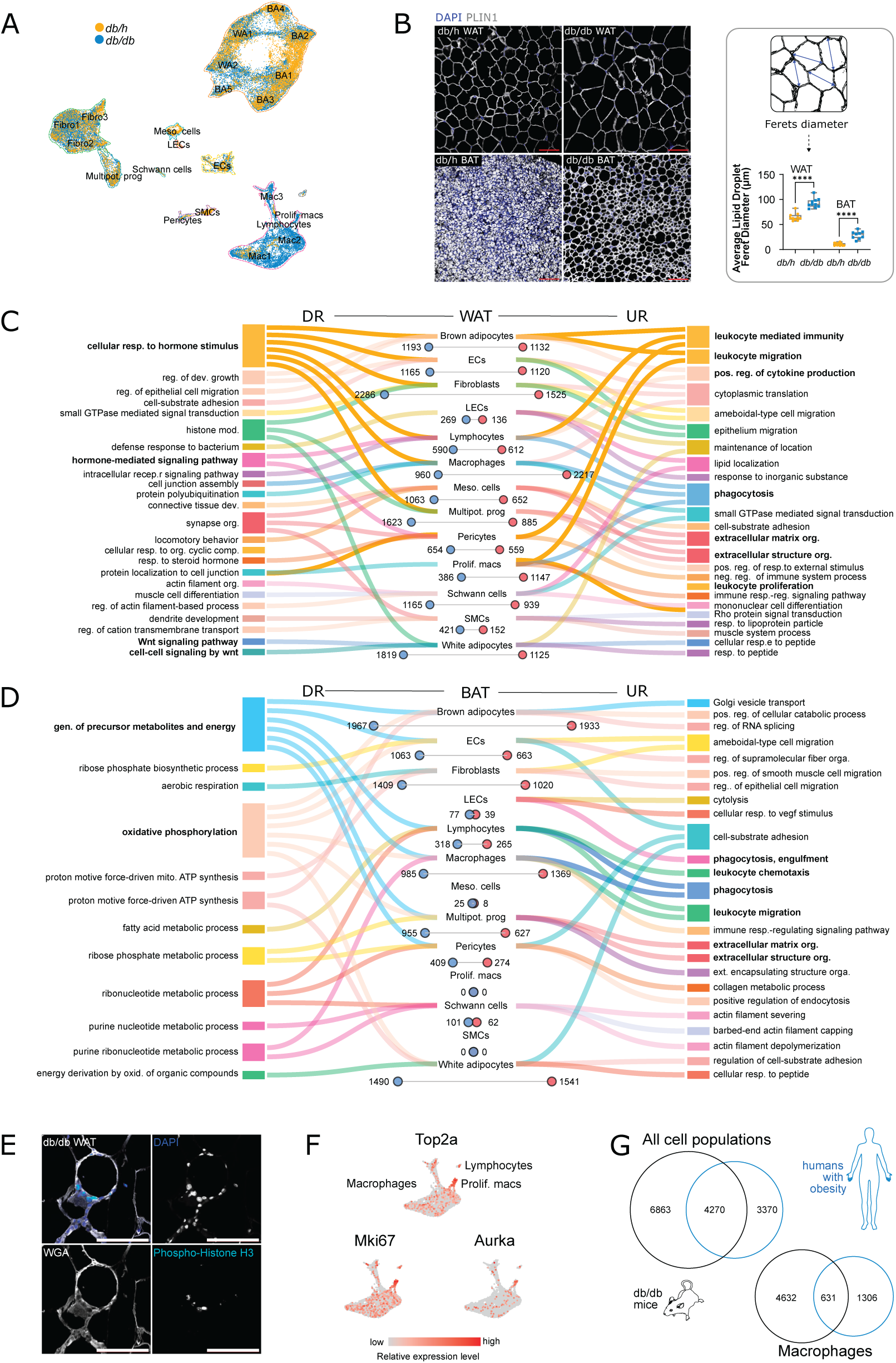
Adipose depot remodeling induced by type-2 diabetes. **(A)** UMAP of both *db/h* nuclei (yellow) and *db/db* nuclei (blue). **(B)** Adipocyte lipid droplet size measured using Feret’s diameter in both *db/h* and *db/db* WAT and BAT. Lipid droplet stained with perilipin 1 (PLIN1; grey) and nuclei stained with DAPI (blue). Corresponding quantification presented. Statistical significance was assumed at P<0.05. ****, *P*<0.0001. **(C)** Sankey plots for WAT demonstrating upregulated (UR; right) and downregulated (DR; left) GO terms and overlayed lollipop plot for associated differentially expressed genes in T2D. **(D)** Sankey plots and lollipop plot for BAT as described previously. **(E)** Immunofluorescence staining of proliferation in crown-like structures in *db/db* WAT through proliferative marker phosphohistone H3 in addition to wheat germ agglutinin (WGA) to mark cell membranes and DAPI to mark nuclei. **(F)** Feature plots showing expression of proliferation markers *Top2a*, *Mki67*, and *Aurka* in macrophage subsets. **(G)** Overlay of differentially expressed genes in all cell populations and macrophages for visceral WAT between the present study and a human obesity study (Emont et al. 2022).

### Type-2 diabetes disrupts cell phenotypes altering adipose tissue homeostasis, energy generation and endocrine functions

Other cell populations that displayed substantially altered gene expression patterns in T2D included white and brown adipocyte subpopulations and fibroblasts (Figure 4C-D). Notable among top GO terms corresponding to upregulated genes in both BAT and WAT across multiple cell populations included *extracellular matrix organization* and *extracellular structure organization* suggesting tissue remodeling. Downregulated terms included *generation of precursor metabolites and energy*, and *oxidative phosphorylation* suggesting decreased energy utilization and possible whitening of classically thermogenic BAT. Genes that were associated with these terms included proteins essential for oxidative phosphorylation such as ATP synthase or cytochrome bc 1 complex (Supplementary Table 3). In WAT, downregulated terms included *cellular response to hormone stimulus*, and *hormone-mediated signaling pathway* indicating a possible dampened hormone response in the tissue. Further, among the top 50 up- and downregulated genes were lysosomal proteases (cathepsins; Cts genes) and leukocyte immunoglobulin-like receptor (LILR) genes including *Lilr4b*, *Pirb* (*Lilrb3*), and *Lilrb4a* with the latter being the only LILR gene in the top 50 differentially expressed of *db/db* BAT (Supplementary Figure 3). Moreover, in *db/db* BAT, the downregulation of heat shock proteins (HSP genes) additionally suggests decreased thermogenesis.

### snRNA-seq derived gene regulatory network inferences in type-2 diabetes

We next sought to determine which GRNs within each depot were driving the changes in adipose tissue cell subsets in disease, focusing on adipocytes, fibroblasts, and macrophages (Figure 5A). Ranking transcription factors based on activity, we found that the top 5 GRNs of these cell types in *db/h* mice were almost always unique across depots. These were altered in *db/db* counterparts and exhibited signatures consistent with the activation of known gene expression programs. For example, Cebpd(+), which is involved in cell apoptosis, proliferation and adipocyte differentiation, was amongst the top-ranked in adipocytes, fibroblasts and macrophage populations in T2D. Additionally, Irf4(+), a critical regulator of lipid handling and of which deficiency induces macrophage-related inflammation^46^, was amongst the top-ranked in BAT *db/h* samples and less so for *db/db* counterparts. Nevertheless, immunohistochemical analyses detected IRF4 in BAT from *db/h* and *db/db* animals (Figure 5B) despite our GRN analysis. These findings suggest possible reduced transcriptional activity and impaired lipid handling in diabetic BAT.

**Figure 5.**
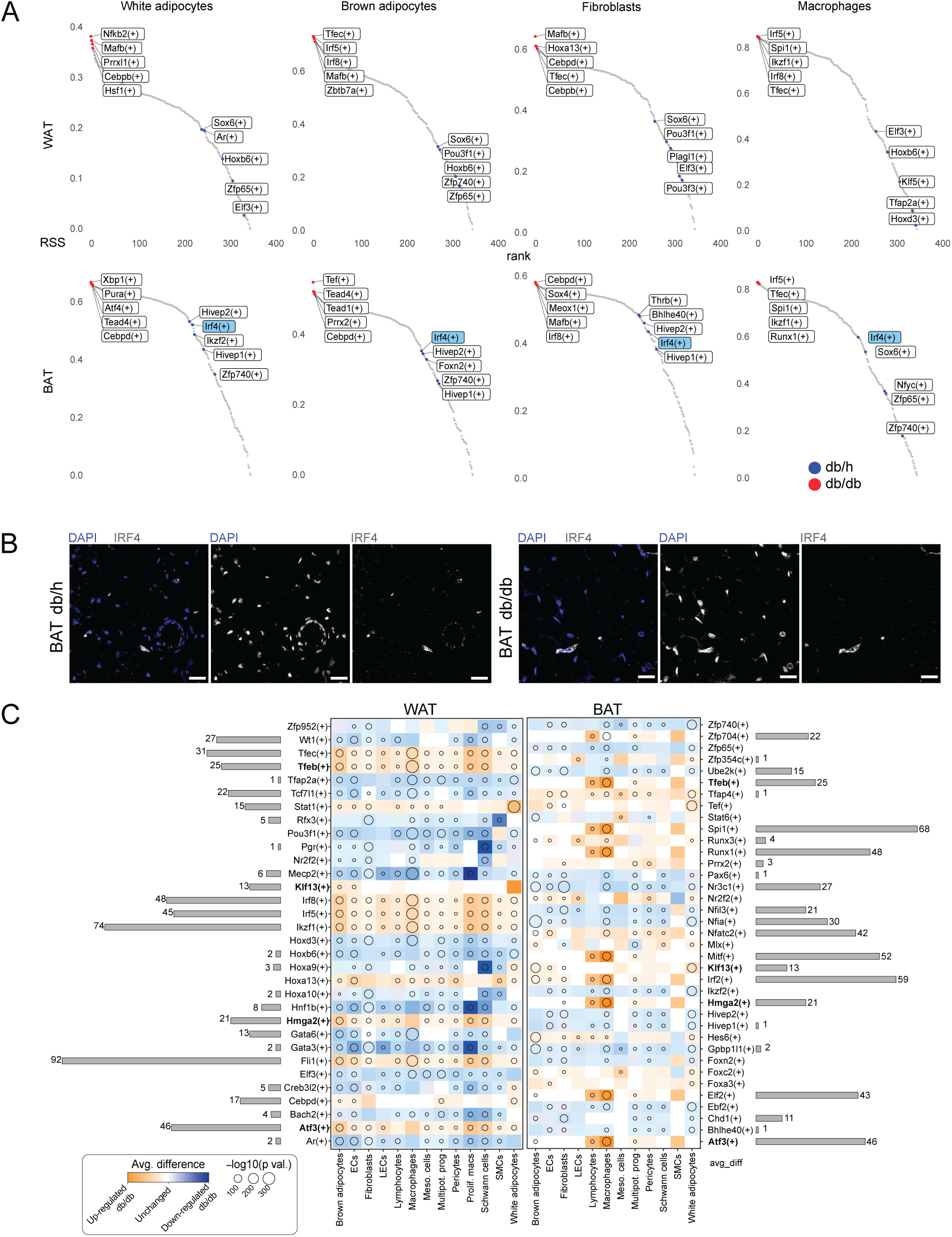
Gene regulatory network alterations in T2D adipose depots and contribution to leukocyte migration. **(A)** RSS plots demonstrating the top 5 activated regulons in *db/db* (red dots) as opposed to *db/h* samples (blue dots) for white adipocytes, brown adipocytes, fibroblasts and macrophages for WAT (top panel) and BAT (lower panel). **(B)** Immunofluorescence validation of transcription factor IRF4 (grey) in both *db/h* and *db/db* BAT tissue with nuclei stained with DAPI (blue). Scale bar represents 25µm. **(C)** Differential enrichment of regulons with upregulated regulons in orange and downregulated regulons in blue. Intensity of color indicates the average expression in that cell population and dot size represents statistical significance. Regulons with bolded text represent those shared across both depots. Bar charts adjacent to regulons indicate the number of target genes within that regulon that are also linked to the GO term *leukocyte migration*.

Next, differentially enriched regulons were assessed. Regulons that showed common changes across both depots included Tfeb(+), Klf13(+), Hmga2(+), and Atf3(+); all being upregulated in a disease context (Figure 5C). Regulons including Tfec(+) and Tfeb(+), among others, showed marked upregulation, particularly in macrophages. TFEB is a transcription factor that plays an essential role in the biogenesis of lysosomal machinery aligning with the upregulated lysosomal protease (Cts gene) expression observed above^47^ (Supplementary Figure 3). To further dissect the influence of transcriptional regulation on the concerted immune response observed (Figure 4), gene targets linked to the GO term *leukocyte migration* were mapped to transcription factors (Figure 5C). Fli1(+), which plays an important role in cell proliferation, activation, differentiation, survival, and the regulation of chemokines^48^, showed the greatest link to this *leukocyte migration* with 92 target genes being associated with this regulon. Other notable regulons included Ikzf1(+) and Spi1(+) which showed 74 and 68 genes linked to this GO term, respectively. Together, these findings highlight the importance of transcriptional regulation for processes, such as inflammation, in T2D adipose depots and highlight possible avenues for targeting TF candidates for therapeutic purposes.

### Biological sex differences in classical adipose depots relating to adipocyte subpopulations

Given the considerable sex differences that exist in obesity and diabetes-associated disorders, we explored potential sex-dependent mechanisms regulating adipose tissue homeostasis and diabetes-induced remodeling. Female and male samples were barcoded using a previously described cholesterol-conjugated oligonucleotide approach^49^ (Supplementary Table 1). Samples were separated downstream based on barcodes. Cells with undetectable barcodes were delineated using sex-specific gene expression (Females: *Xist*; Males: *Ddx3y*, *Eif2s3y*, *Gm29650*, *Kdm5d*, *Uty*)^22^.

Comparison of cells from females and males across each depot uncovered extensive sex-specific differences. Sex-specific gene expression analysis found that within WAT, mesothelial cells, fibroblasts, macrophages and white adipocytes exhibited sex-specific gene expression (Figure 6A). In BAT, gene expression variation was primarily within adipocyte subpopulations. When the cellular composition of these depots was considered, females and males had differing proportions of adipocyte subpopulations (Figure 6B). Specifically, *db/h* female BAT had high proportions of BA1 and BA2 cells, while *db/h* male BAT had higher proportions of BA3 and BA4. Assessment of key genes involved in thermogenesis showed higher expression levels in female *db/h* BAT while these dimorphisms were not as evident in a diabetic context (Figure 6C). Analysis of the top 5 GO terms for brown adipocytes in *db/h* BAT provided some insight into biological pathway differences showing female upregulation of *adaptive thermogenesis*, *cellular response to hormone stimulus* and *fatty acid metabolic process* in females, and upregulation of *protein folding*, *oxidative phosphorylation*, *ATP metabolic process,* and *proton motive force-driven ATP synthesis* pathways in males (Figure 6D). Genes linked to these pathways encoded ATPase subunits, and heat shock proteins in males while in females they encoded thermogenesis-related genes, including *Ppargc1a* and *Ucp1*, alongside *insulin receptor substrate 1* (*Irs1*) (Supplementary Tables 4 and 5). These findings align with the long recognized increased thermogenic capacity of female BAT driven by mitochondrial size, abundance and UCP1 content^50–53^.

**Figure 6.**
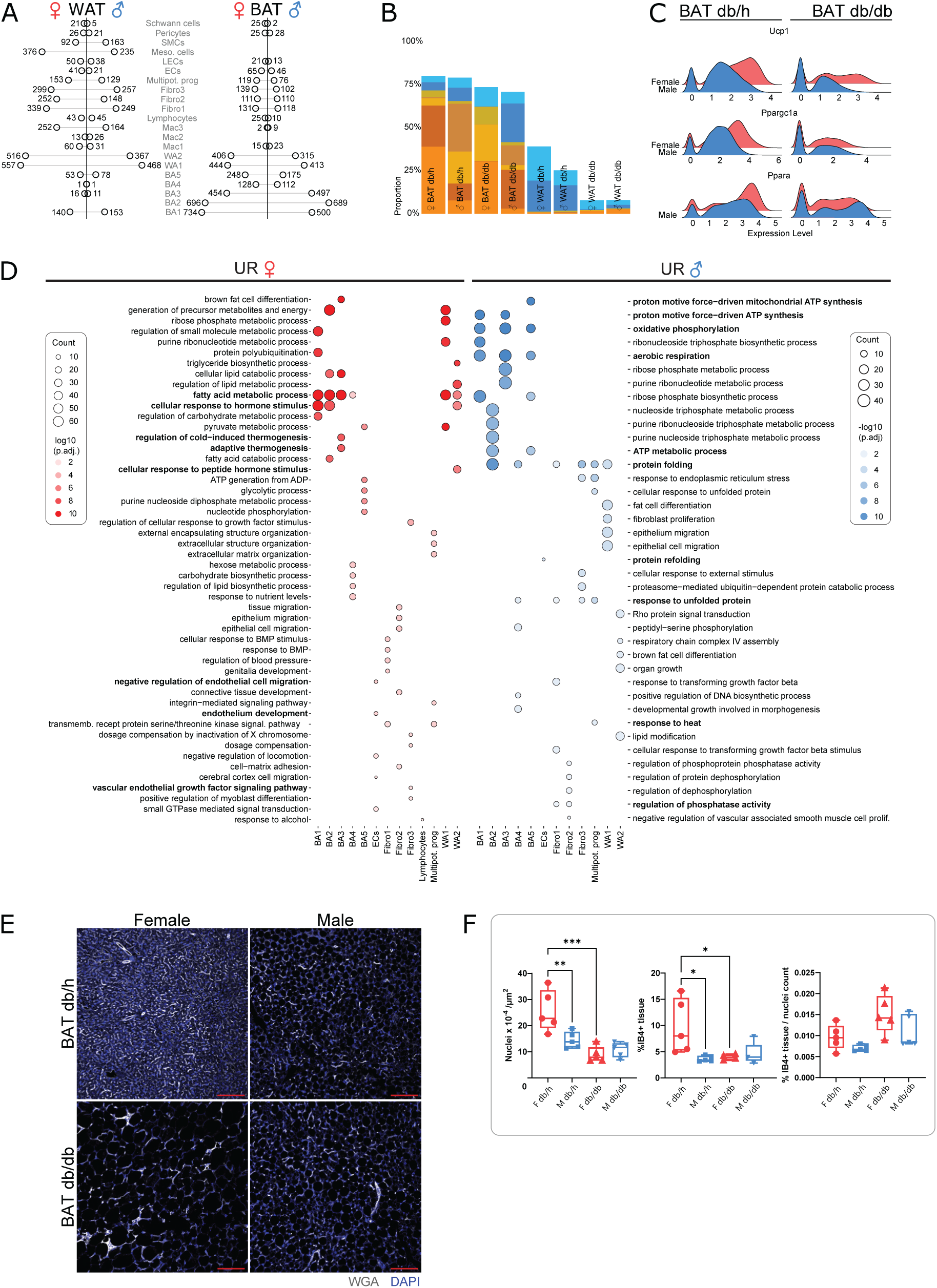
Sex differences in gene expression and tissue architecture across adipose depots. **(A)** Lollipop plot demonstrating sex differences in gene expression between females and males in *db/h* WAT and BAT. **(B)** Compositional bar chart demonstrating the contribution of adipose tissue cell populations to each depot and how they differ between conditions and biological sexes. **(C)** Ridgeplots showing female and male sample expression of key thermogenic genes between *db/h* and *db/db* conditions in BAT. **(D)** Top 5 gene ontology terms for each cell population where dot size represents the number of genes linked to this GO term and dot color represents the statistical significance. GO terms of interest are highlighted in bold. **(E)** Immunofluorescence staining of BAT vascularization using isolectin B4 (IB4; red) and nuclei stained with DAPI (grey) demonstrating differences in tissue architecture between females and males in both *db/h* and *db/db* conditions. Scale bar represents 100µm. **(F)** Quantification of vascularization (% of IB4+ tissue area) and cell density (number of DAPI+ nuclei) histology with female data points presented in pink and males in blue. Statistical significance was assumed at P<0.05. * *P*<0.05, ** *P*<0.01, *** *P*<0.001.

### Higher cell density and vascularization in female BAT

Finally, as many endothelial-related GO terms were noted in the analysis for female BAT we sought to assess vascular differences between sexes (Figure 6D). Efficient blood flow to the tissue is essential for the supply of free fatty acids for energy metabolism and generation. Differences between biological sexes have previously been observed in gonadal WAT^54,55^ yet, to date, no sex differences in BAT vascular networks have been characterized. Staining of female and male *db/h* BAT showed more cells and IB4+ vasculature in females (*P*=0.0091 and *P*=0.0206, respectively; Figure 6E-F). However, normalization of IB4 staining to the number of nuclei revealed that the ratios were similar. These data indicate that, while female tissue may be more densely packed, vascularization-to-cell ratios remain comparable. Strikingly, cell and vascularization density sex differences were lost in T2D BAT alongside reduced gene expression differences across adipocyte populations (Figure 6E-F). Together, these data highlight that BAT sex differences that exist in a healthy state may be dampened or lost in disease.

## Discussion

Pathogenic adipose tissue expansion is a well-established risk factor for cardiometabolic disease. Recent research has been directed towards understanding how the deterioration of adipose tissue can contribute to disease in both an obese and high-fat diet context at a single-cell level^14–21^. Limited T2D-focussed human work has characterized insulin-regulating adipocyte subpopulations and molecularly distinct preadipocyte populations associated with diabetes^7,13,21^. However, despite this, we are still lacking a precise understanding of the cellular and molecular underpinnings of how these tissues change in the context of T2D. To address this, we conducted snRNA-seq on classical adipose depots including gonadal WAT and interscapular BAT from both *db/h* and *db/db* female and male mice.

Characterization of these depots using snRNA-seq showed that while similarity in cellular heterogeneity existed between depots, the contribution of various cell subsets to the overall tissue cellularity differed. Moreover, gene expression patterns differed across WAT and BAT highlighting the distinct functions of these tissues. WAT is classically recognized for its lipid-storing functions, while BAT is known for its heat-producing and energy-expending role. Analysis of adipocyte subtypes highlighted that cell populations have unique functions across depots with BAT populations upregulating metabolic processes and WAT populations upregulating WNT-related pathways. WNT signaling has important adipose-specific functions including regulating lipogenesis, inhibiting adipogenesis^56^, and the modulation of structural WNT-regulated adipose tissue-resident cells, termed SWAT cells, which can move toward both adipogenic or progenitor-like states^57^. Finally, the assessment of GRN inferences validated the observed depot-specific gene expression patterns.

Our analysis of non-diabetic and T2D mice also revealed several features of pathogenic remodeling of adipose. These include an increase in macrophage abundance across both WAT and BAT, consistent with observations by others^14,16–21^. Our findings align with the more recent recognition of proliferative macrophage importance in CLS formation^44,45,58,59^ and highlight their presence in a model of T2D. Assessment of GRN dynamics in T2D also showed a notable increase in TFEB transcription factor activity in macrophage subsets. TFEB is an important transcription factor in the biogenesis of lysosomal machinery^47,60^. Lysosomal degradation of hypertrophic adipocytes is a vital function of macrophage CLSs where these phagocytic cells secrete lysosomal enzymes to create an environment wherein dying adipocytes are degraded through a process known as exophagy. Lysosomal proteases are vital for this process with CTSB being specifically identified as key for perilipin 1 (PLIN1; a lipid-droplet coating protein) degradation^61–63^. Indeed, these lysosomal proteases are upregulated in *db/db* samples (most predominantly in macrophages) which was further corroborated through Tfeb(+) expression –TFEB being a known regulator of Cts genes^47^.

In line with the noted adipocyte lipid droplet expansion, BAT showed downregulation of energy generation in T2D. BAT whitening has gathered much attention in the literature due to the hijacking of BAT-driven energy expenditure to improve metabolic health. A wealth of research has been directed into exploring therapeutic options to increase this expenditure including the use of cold exposure, exercise, and pharmacological and dietary interventions^6^. Assessment of genes showing decreased expression in T2D included heat shock proteins. Moreover, GO analysis of downregulated genes showed enrichment of genes related to vital oxidative phosphorylation complexes, further supporting the notion of BAT moving into a lipid-storing state.

Finally, previous research has highlighted biological sex differences between adipose depots, including functionality differences in BAT. Female BAT has greater mitochondrial abundance and size, in addition to higher mRNA and protein levels of UCP1 and other thermogenic genes^50–53,64–66^, which aligns with our data. Further, although single-cell datasets have shown brown adipocyte populations with differing heat-producing capacities^10,67,68^, sex differences have not been characterized. Here we found that differences exist in adipocyte subpopulation abundance across sexes. Moreover, histological analysis showed greater cellular and vascular density in female BAT, with differences only being identified in gonadal WAT in the past^54,55^. VEGFA is long linked to BAT activity, with *Vegfa* deletion driving BAT whitening resulting in decreased expression of thermogenic gene *Ucp1* and *Ppargc1a*^69^. Moreover, estrogen receptor 1 is known to regulate VEGFA in adipose tissue possibly driving the sexual dimorphism^70^. However, T2D appeared to abrogate these sex differences in vascular and gene expression.

While this study offers new insights and a resource for studying adipose, several limitations require noting. First, the model of T2D used in this study (*db/db*) is a leptin receptor mutant meaning changes in depots may be driven by the loss of this signaling rather than the diabetic phenotype. While we compared our data to that of humans with obesity, data from this study should be further compared across other models of T2D. Second, it is important to note that *db/db* mice are characterized by obesity, thus findings in the study may not be strictly attributed to T2D alone. It has been shown, however, that *db/db* mice appear more ‘diabetic’ than *ob/ob* mice and have less liver failure^71^. Third, mouse adipose depots, particularly BAT, inconsistently recapitulate the human context. Thermoneutral BAT is more similar to human BAT and could be used in future studies, particularly those considering biological sex differences^72,73^. Finally, as this dataset was produced using nuclei, cytosolic mRNA is missing and therefore important transcriptomic information could be absent from our dataset.

Overall, this study offers a unique insight towards adipose tissue biology and the multifaceted mechanisms driving its maladaptive remodeling in T2D. It also informs our understanding of the impact of biological sex in adipose homeostasis and disease. In addition to the single-cell transcriptomic resource presented here, further analysis of different models, time points and genetic backgrounds will advance our knowledge of adipose remodeling, potentially pointing to mechanisms that may be therapeutically manipulated to address tissue dysfunction induced by metabolic disorders such as T2D.

## Research design and methods

### Animals, physiological assessment, and tissue isolation

All animal-related experiments were approved by the Alfred Research Alliance (ARA) Animal Ethics Committee (Ethics number: E/1990/2020/B) and were performed in accordance with the National Health and Medical Research Council of Australia. 20-week-old female and male B6.BKS(D)-Lepr^db/J^ (*db/db;* referred to as T2D) (strain #:000697, The Jackson Laboratory) or heterozygous littermates (*db/h;* referred to as ND) in the C57BL/6J background^74–76^ were housed under 12h light/dark cycles at 22°C with access to both food and water *ad libitum*. Homozygous *db/db* mice are a well-established model of T2D, having a spontaneous autosomal recessive global knockout for the leptin receptor. At 19-weeks of age, whole-body composition analysis was performed on mice using Echo-MRI™ 4-in-1 700 Analyser (EchoMRI, Houston, TX, USA). Long-term blood glucose levels were measured via HbA_1c_ at endpoint (Cobas b 101 POC system, Roche Diagnostics). At 20 weeks of age, mice were administered an intraperitoneal injection of heparin (100 units per mouse) 20-30 minutes prior to euthanasia using CO_2_ inhalation, which was followed by subsequent cardiac exsanguination. As described previously^77^, the thoracic cavity was exposed, and the right atrium was cut. The pericardium was removed, and the inferior vena cava was cut. The apex of the left ventricle was pierced using a 30-gauge hypodermic needle for perfusion with buffers specified below.

### Stromal Vascular Fraction Flow Cytometry Processing

Mice were perfused with cold ethylenediaminetetraacetic acid (EDTA) at a rate of 2 mL/min/mouse before gonadal white adipose and interscapular white and brown adipose depots were excised and snap frozen. Tissues were washed briefly in clean PBS with 0.9 mM CaCl_2_ (C4901, Sigma) before being minced using curved scissors into ∼2 mm pieces in a petri dish and transferred into 3 mL of enzymatic solution (2 mg/mL collagenase IV (LS004188, Worthington Biochem) and 1 mg/mL dispase II (04942078001, Roche) in PBS with 0.9 mM CaCl_2_). Samples were triturated three times in 15-minute intervals using a P1000 pipette for a total of 45 minutes then centrifuged at 400xg for 4 minutes at 4°C. The floating fraction was then carefully removed using a P1000 pipette and discarded. Using the remaining supernatant, the pellet was resuspended before being filtered through a 70 μm filter (43-50070-50, pluriSelect) into a tube containing PBS with 0.9 mM CaCl_2_. The sample was pulled through the filter twice, followed by the addition of count beads (C36950, Invitrogen) before centrifugation at 200xg for 15 minutes at 4°C - no breaks - for debris clearance. The supernatant was removed carefully, and the pellet was resuspended in Fx buffer (1XHBSS (14185-052, Gibco), 2% FCS in ddH2O (10977015, Invitrogen)) before centrifugation at 400xg for 4 minutes at 4°C. The supernatant was removed, and the pellet was resuspended in 50 μL of Fx buffer. A volume of 43μL for each sample was transferred to 1.5 mL tubes containing Fc block (101320, BioLegend) on ice for 5 minutes and the remaining samples were pooled and used for single color controls. Samples were then incubated in tubes containing antibody mix (Supplementary Table 1) on ice for 20 minutes covered from light. Calcein Blue (1:500; 65-0855-39, Invitrogen) was added to each sample and incubated at 37°C for 10 minutes. Samples were washed using Fx buffer before centrifugation at 400xg for 4 minutes at 4°C. Supernatant was removed and samples were resuspended in Sytox Green (1:1000; S34860, Invitrogen) in Fx buffer before being filtered into FACS tubes (38030, StemCell) and analyzed on a BD Fortessa II flow cytometer.

## Statistical Analysis

FlowJo™ V10.8 Software (for Windows BD Life Sciences) was utilized to compensate and analyze flow cytometry data. A detailed gating strategy is outlined in Supplementary Figure 1C. For sex-independent analyses within each depot, raw data were normalized to the average value for all *db/h* samples for that batch. For sex-dependent analyses, the prior was conducted with only female samples for each batch. GraphPad Prism v9.2.0 (GraphPad Software San Diego, California USA, www.graphpad.com) was subsequently used to analyze data and conduct statistical analyses. Outliers were excluded using GraphPad Prism’s inbuilt ROUT outlier identification method (Q=1%) on data within each cell population. For between depot analysis, data were treated as non-parametric and analyzed using a Kruskal-Wallis test. For within depot analysis of T2D changes and sex differences, data were treated as non-parametric and analyzed using a Mann-Whitney test. *P* values less than 0.05 were considered statistically significant.

### Nuclei extraction

Mice were perfused with cold ethylenediaminetetraacetic acid (EDTA) at a rate of 2 mL/min/mouse before gonadal white adipose and interscapular white and brown adipose depots were excised and snap frozen. Frozen adipose tissue samples (ranging from ∼25 to 34.7 mg for WAT and ∼60.9 to 65.5 mg for BAT) were mechanically homogenized in a previously published^78^ nuclei purification buffer (NPB) (containing 5% bovine serum albumin, 0.2% Nonidet^TM^ P 40 Substitute (74385, Sigma-Aldrich), 1 mM dithiothreitol (D0632, Sigma-Aldrich), 1X cOmplete^TM^ EDTA-free Protease Inhibitor Cocktail (11873580001, Roche), 0.2 U/µL of Protector RNase Inhibitor (3335399001, Roche) in Dulbecco’s PBS (14190-250, Gibco)) with a 5 mm diameter stainless steel ball (Qiagen, 69989) in Eppendorf Safe-Lock LoBind tubes (0030120.094, Eppendorf) using a Qiagen TissueLyser II. BAT and WAT were homogenized at 30 Hz for 30 and 50 seconds, respectively, in 10-second intervals (Supplementary Figure 1A). Samples were placed on ice post-homogenization and incubated for 15 minutes. Homogenized samples were then pulled through 20 µM pluriSelect Cell Strainers (43-50020-03, pluriSelect) to remove debris and centrifuged at 1000xg for 5 minutes at 4°C. Pellets were then resuspended in cold nuclear Fx Buffer (containing 1X HBSS, 2% fetal calf serum, 0.2 U/µL of Protector RNase Inhibitor). Nuclei were barcoded using cholesterol modified barcodes and anchors as previously described^49^. Briefly, nuclei were first incubated with the anchor-barcode mix at a final concentration of 200nM for 5 minutes on ice before being further incubated with a co-anchor at a final concentration of 200 nM for 20 minutes (Barcodes in Supplementary Table 1). Samples were twice washed in cold nuclear Fx buffer before samples within each group were pooled and stained with DAPI (1 µg/mL; D1306, Invitrogen) in cold nuclear Fx buffer. Nuclei were filtered through a 35 µM filter (38030, StemCell) before being sorted on the BD FacsARIA Fusion to remove debris (Alfred Research Alliance FlowCore).

### Single-nucleus transcriptomic library preparation and sequencing

FACS isolated nuclei were counted using a hemocytometer and pooled as detailed in Supplementary Table 1 for single nucleus transcriptomic library preparation using the Chromium controller (10X Genomics). Approximately 40,000 nuclei were loaded into each channel and processed using the Chromium Single Cell 3’ v3.1 reagent kit (10X Genomics). After capture and lysis, cDNA was synthesized and amplified for 12 cycles following the manufacturer’s protocol. Once amplified, the cDNA was used for downstream sequencing on four lanes of an Illumina NovaSeq 6000 to an approximate depth of 50,000 reads and 3000 genes per nucleus and a sequencing saturation of around 50-61.5% (see Supplementary Table 1).

### Analysis of single-nucleus RNA sequencing data

Raw sequencing data in FASTQ format was processed using Cell Ranger v6.1.1 (10X Genomics) with the inclusion of intronic reads before downstream analysis. Reads that were confidently mapped to the genome were aligned to the *Mus musculus* mm10 transcriptome before the expression of each transcript was quantified for each nucleus. Analysis was carried out using R v4.1 and v4.2 and Seurat v4.1.1 (satijalab.org/seurat). Nuclei with <200 and >6500 and >7000 unique genes expressed for GEX_A (BAT_dbh) and GEX_B (BAT_dbdb), respectively, and nuclei having <200 and >7500 unique genes expressed were excluded for GEX_C (WAT_dbh) and GEX_D (WAT_dbdb), respectively. Nuclei with a percentage of reads mapping to mitochondrial genes greater than 2, 1, 1, and 0.5 standard deviations were removed from GEX_A, GEX_B, GEX_C and GEX_D, respectively, to keep the top 80% of nuclei for analysis. SCTransform^79^ was used to normalize the data and regress out ribosomal protein genes (genes starting with Rps and Rpl) and mitochondrial genes to reduce technical artifacts. Four objects were created for each sequencing run and nuclei were clustered. Low quality clusters were removed based on nFeature_RNA, nCount_RNA and percent.mt. Remaining cells were re-clustered, and doublets were removed using DoubletFinder^80^ prior to sample integration. Standard Seurat integration approach was performed with 4000 integration features used and canonical correlation analysis (CCA) was utilized to find the integration anchors. Sex-based separation was primarily conducted using the barcoding approach described above. Sex barcodes were demultiplexed using MULTIseqDeemux() function in Seurat. Nuclei that were predicted as doublets were either further separated based on our previously published approach^22^ that considers the expression of sex-specific genes (*Xist*, females; *Ddx3y*, *Eif2s3y*, *Gm29650*, *Kdm5d*, *Uty*, males) or discarded from analysis if they showed co-expression. Sex information then was embedded into the integrated Seurat object. Dimensionality reduction was conducted using 40 principal components and a resolution of 0.8 to explain variability not expected by chance to cluster the nuclei into cell populations. Uniform Maximum Approximation and Projection (UMAP) was used to visualize the data in two dimensions and ‘FindAllMarkers’ was used to identify cell clusters. As observed in previous adipose single cell datasets, contaminating epididymal cells, ovarian epithelial cells and skeletal muscle cells were removed from analysis. The final dataset included 51,877 nuclei across 28 samples (7 samples in each of the 4 groups) comprising 22,526 nuclei in WAT and 29,351 nuclei in BAT from db/h and db/db samples.

### Differential expression analysis

To assess differentially expressed (DE) genes between conditions and sexes, ‘MASTcpmDetRate’ was used. MASTcpmDetRate is a modified version of ‘MAST’^81^ which considers cellular detection rate and has been successfully used in the past for benchmarking experiments that deemed this approach superior to others^82^. First, ribosomal RNA was removed from the dataset and DE analysis was conducted. Genes with an uncorrected *P* value less than 0.01 were considered statistically significant and, thus, differentially expressed between groups. Data was primarily visualized using the ggplot2 package.

### Gene ontology analysis

Gene ontology (GO) enrichment analysis of the above DE genes was performed using the ‘enrichGO’ function from the clusterProfiler package (v4.4.4)^83^. DE genes were mapped to genomes from geneontology.org. Enrichment analysis of GO Biological Process terms (GO terms) was calculated using all genes identified in the dataset as the background gene set (with minimum and maximum gene set sizes as 10 and 500, respectively). The ‘simplify’ function was used to calculate similarities between GO terms. GO terms with a similarity higher than the 0.7 cut-off were discarded from analysis. Terms with a *P* value less than 0.05 were considered statistically significant GO terms. Data was visualized using the ggplot2 package.

### Gene expression overlap analysis

To determine the similarity of differentially expressed genes between mouse *db/db* and human obesity datasets, a publicly available scRNA-seq dataset^21^ of visceral WAT and present data on gonadal (visceral) WAT samples from *db/db* mice were compared. For the human obesity dataset, samples with a BMI greater than 30 were classed as ‘obese’ while those less than 25 were classed as ‘lean’. DE analysis was conducted between control and treatment groups (lean vs. obese for humans and *db/h* vs. d*b/db* for mice), as described above. Human gene orthologs were converted to mouse orthologs using the AnnotationDbi package (v1.58.0). DE genes with a *P* value less than 0.01 were isolated from each dataset and the ‘overlap’ function within RVenn (v1.1.0) was used to determine both common and unique DE genes across datasets. The number of common and unique genes were visualized using RVenn (v1.1.0), Euller (v6.1.1), VennDiagram (v1.7.3) and Vennueler (v1.1.3).

### Tissue preparation for histological validation

Age and sex-matched animals were euthanized using CO2 inhalation cardiac exsanguination. Animals were then perfused with PBS with 200mM KCl (P9541, Sigma-Aldrich) at a rate of 2 mL/min/mouse until the liver cleared of blood. Mice were then perfused with 4% PFA (P6148-500G, Sigma) in PBS with 200mM KCl at a rate of 1 mL/min/mouse. Tissue was kept in 4% PFA in PBS overnight before being stored in PBS with PBS with 0.09% NaN_3_ (S2002-25G, Sigma) at 4°C short-term. Tissue was embedded in paraffin by the Monash Histology Platform (Monash University, Clayton, VIC, Australia).

### Tissue section staining

FFPE blocks were cut on a microtome (RM 2235, Leica Biosystems) at 10 µm on SuperFrost plus slides then dried overnight at 37°C. Paraffin was melted at 80°C for 1 minute before slides were washed three times with fresh xylene for 10 minutes each (534056-4L, Honeywell). Slides were rehydrated in 100% (v/v) EtOH, 100% (v/v) EtOH, 90% (v/v) EtOH, 70% (v/v) EtOH and 50% (v/v) EtOH for 5 minutes each before being submerged in ddH_2_O for 1 minute. Slides were washed twice in Tris-EDTA, pH 9.0 (10 mM Tris (BIO3094T-1KG, Astral Scientific), 1 mM EDTA (EDS-500G, Sigma) in ddH2O) before being microwaved for 15 minutes in 1 L of Tris-EDTA, pH 9.0. Slides were washed twice with PBS-T (PBS, 0.05% Tween-20) before being permeabilized for 10 mins in 0.5% Triton X-100 (T9284, Sigma) in PBS. Slides were washed twice with PBS for 5 minutes each before blocking with goat block (2% goat serum (G9023-10ML, Sigma), 1% BSA (A3608-100g, Sigma), 0.1% cold fish skin (G7765-250ML, Sigma), 0.1% Triton X-100 (T9284, Sigma) for 1 hour at room temp. Slides were then incubated with primary antibody in goat block (PLIN1, 1:200, AB3526, Abcam; Isolectin B4/IB4, 1:50, L2140, Sigma-Aldrich; Interferon Regulatory Factor 4/IRF4, 1:200, PA5-115603, Invitrogen) overnight at 4°C in a humidified chamber. Sections were washed three times in PBS-T for 5 minutes each. Slides were incubated with secondary antibody in PBS-T for 1 hour at room temp. Sections were then washed three times with PBS-T with AF488 conjugated Wheat Germ Agglutinin (WGA) (29022, Jomar Life Research) for 5 minutes before a final wash with DAPI to stain nuclei in PBS-T for 10 minutes. Sections were mounted under a coverslip before imaging using a 20X multi-immersion objective and glycerol on a Nikon A1R confocal laser scanning microscope.

### Statistical analysis of quantified data

For immunofluorescence-based validation, micrographs were analyzed in Fiji image analysis software^84^. Macros were utilized for IB4, nuclei and adipocyte lipid size quantification. Briefly, for tissue sections stained with IB4 and DAPI, the total tissue area and IB4 stained tissue area (determined using thresholding) were measured within a field of view. The percentage of IB4-positive tissue space was calculated. Nuclei (DAPI-positive) were selected with thresholding, quantified, and divided by the total area to give the number of nuclei per µm^2^. Finally, thresholding of PLIN1-positive lipid droplets facilitated the subsequent measurement of Feret’s diameter (µm) for each PLIN1-positive lipid droplet. Calculations using raw data from Fiji were conducted in Microsoft Excel before processed data were transposed into GraphPad Prism (v9.2.0). GraphPad Prism (v9.2.0) was used to conduct unpaired t-tests on comparisons between two groups and ordinary one-way ANOVAs on comparisons between three or more groups. Statistical significance was determined where *P*<0.05.

## Acknowledgments

We would like to acknowledge the Boon Wurrung and Woi Wurrung (Wurundjeri) Peoples of the Kulin Nation on which our research was formulated, conducted, and reported on. We would like to pay our respects to their Elders past, present and emerging.

The authors acknowledge use of the facilities at the AMREP Flow Cytometry Platform, the Monash Histology Platform and the Baker Heart and Diabetes Institute Microscopy Platform.

## Author contributions

A.R.P. supervised the study. A.R.P., G.E.F., C.K., T.L.G., and C.D.C. conceived and designed the experiments and collected the data. I.H., M.S.I.D., T.L.G., C.K., C.D.C., and A.R.P. performed the data analysis with I.H. and M.S.I.D. leading the computational analysis. A.R.P. and T.L.G. wrote the manuscript. A.R.P., B.G.D., T.L.G., G.E.F., C.K., C.D.C., I.H., B.G.D and M.S.I.D. reviewed, edited, and approved the manuscript.

## Ethics approval

All animal-related experiments were approved by the Alfred Research Alliance (ARA) Animal Ethics Committee (Ethics number: E/1990/2020/B) and were in accordance with NHMRC guidelines.

## Data availability

All raw and processed sequencing data, and corresponding scripts, will be made available upon publication.

---

**Supplementary Figure 1.**
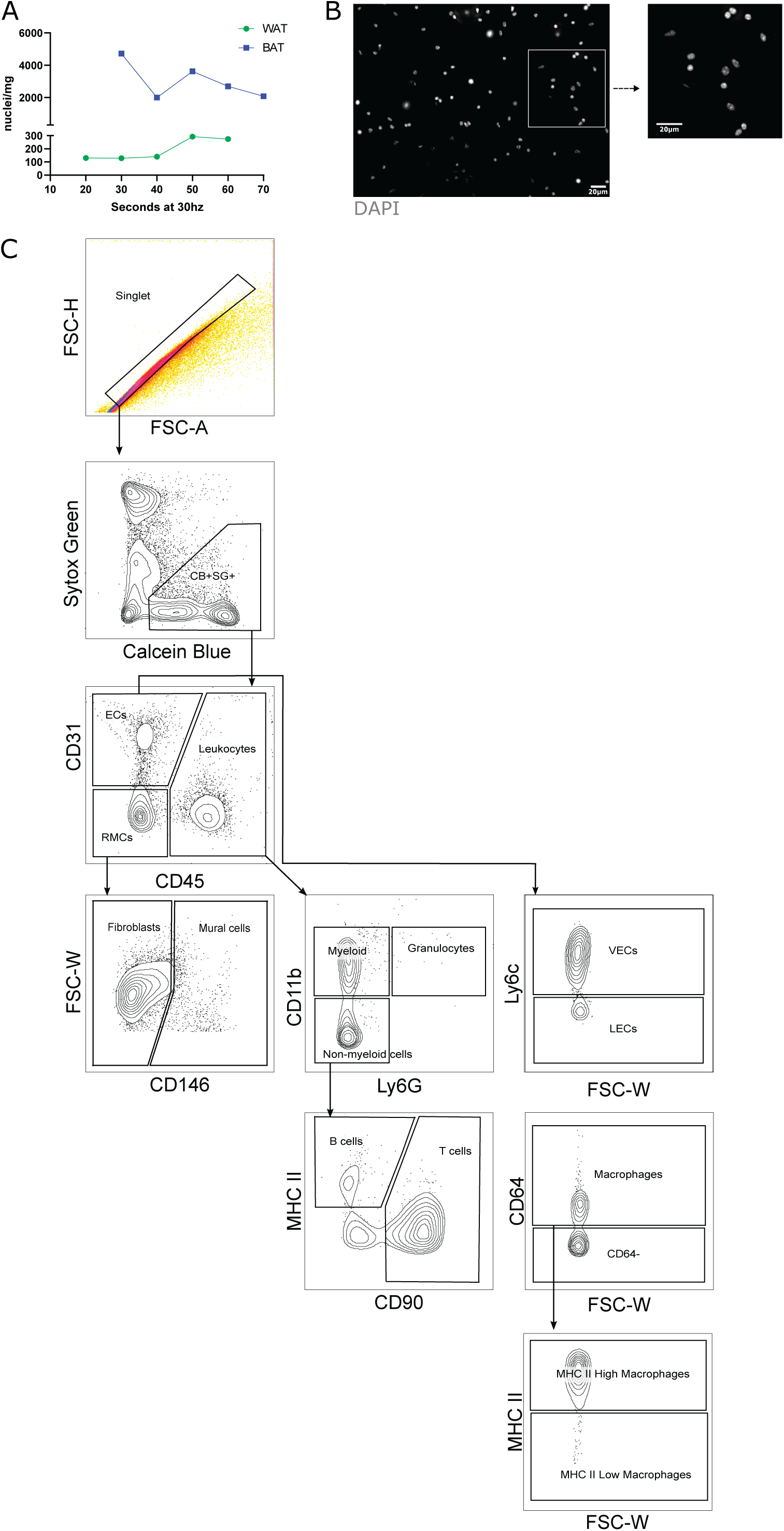
Optimization of nucleus isolation and flow cytometry gating strategy. **(A)** Flow cytometry-based optimization of the time length required to mechanically digested at 30hz using the Qiagen Tissue Lyser to get optimal nuclei/mg of tissue yield. **(B)** micrographs showing nuclei post Qiagen digestion and fluorescence-activated cell sorting to remove cellular debris. Scale bar represents 20µm. **(C)** Flow cytometry gating strategy for SVF cell population analysis. Viable and metabolically active single cells were gated, and cell populations were delineated based on staining by antibodies specific for cell surface markers. WAT, white adipose tissue; BAT, brown adipose tissue; FSC-H, forward scatter height; FSW-A, forward scatter area; FSC, forward scatter width; CB, calcein blue; SG, sytox green; ECs, endothelial cells; RMCs, resident mesenchymal cells; VECs, vascular endothelial cells; LECs, lymphatic endothelial cells.

**Supplementary Figure 2.**
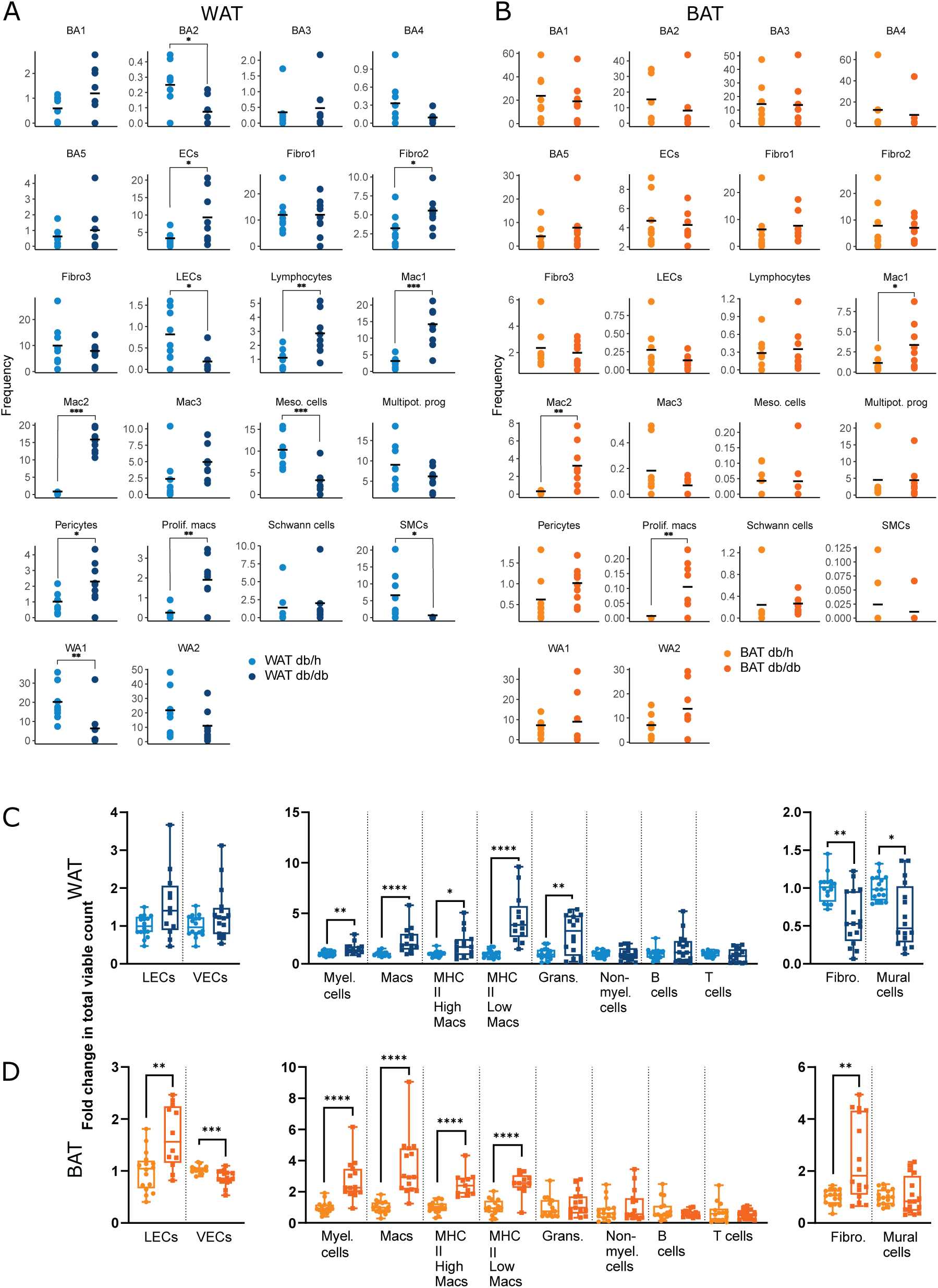
Cell population abundance changes in T2D. **(A)** Cell proportion differences between *db/h* and *db/db* samples based on nuclei count in WAT and BAT. Y axis indicates frequency of total cell count. **(B)** Contribution of adipose tissue cell populations to flow cytometry total viable cell count with differences between *db/h* (light color) and *db/db* samples (dark color) being shown in WAT (blue) and BAT (orange) depots. BA, brown adipocyte; WA, white adipocyte; ECs, endothelial cells; Meso. cells, mesothelial cells; Multipot. prog, multipotent progenitors; Prolif. macs, proliferating macrophages; SMCs, smooth muscle cells; WA, white adipocyte; LECs, lymphatic endothelial cells; VECs, vascular endothelial cells; Myel. cells, myeloid cells; Macs, macrophages; Grans, granulocytes; Non-myel. cells, on-myeloid cells; Fibro, fibroblasts. Statistical significance was assumed at P<0.05. * *P*<0.05, ** *P*<0.01, *** *P*<0.001, **** *P*<0.0001.

**Supplementary Figure 3.**
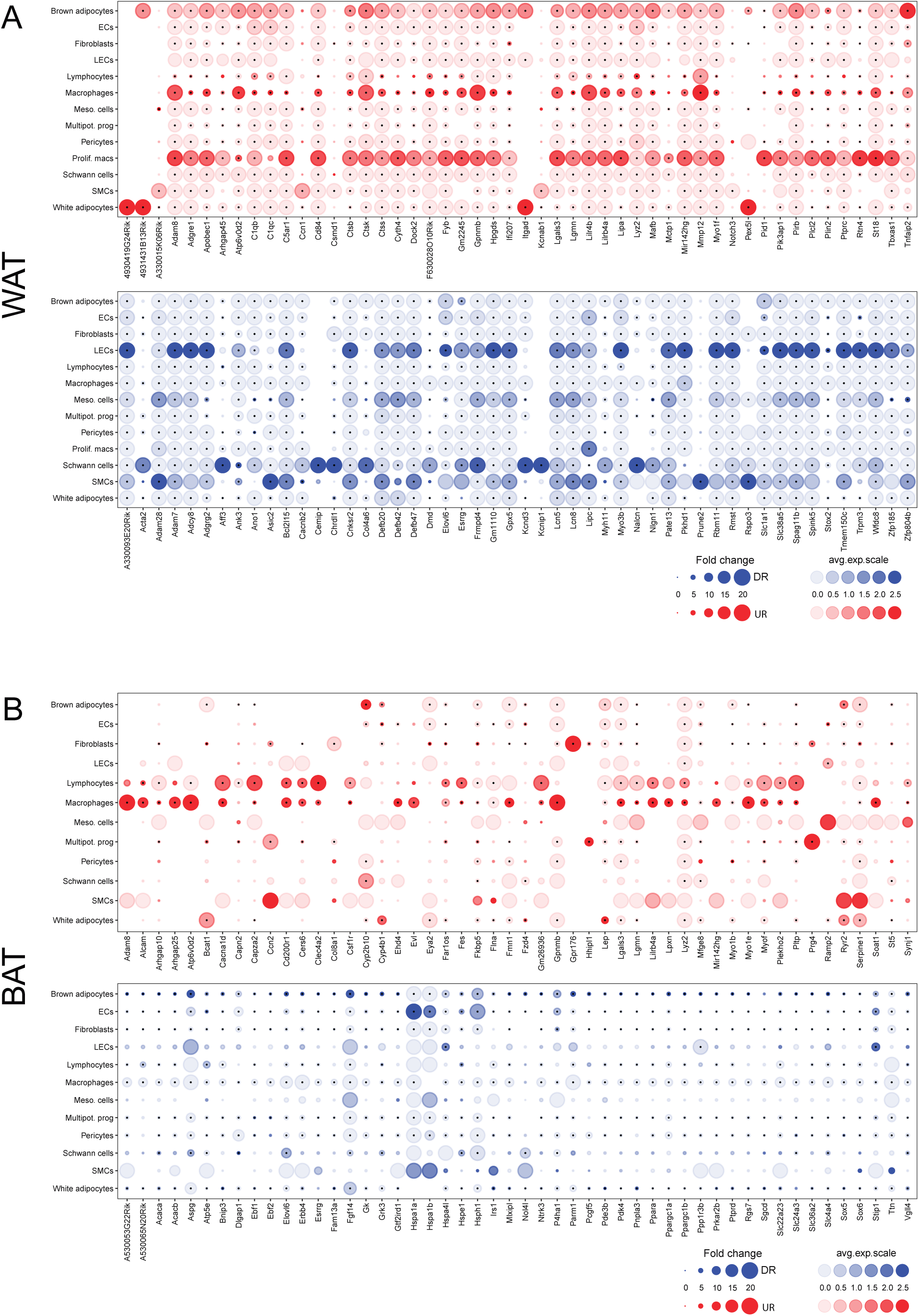
Top 50 upregulated and downregulated genes in T2D WAT and BAT. Genes up-(red) downregulated (blue) between *db/h* and *db/db* cell populations in WAT **(A)** and BAT **(B)**. Dot size represents the fold change (FC) difference between conditions while color intensity indicates the expression level (avg. exp. scale). ECs, endothelial cells; LECs, lymphatic endothelial cells; Meso. cells, mesothelial cells; Multipot. prog, multipotent progenitors; SMCs, smooth muscle cells.

## References

1. Oikonomou, E.K., and Antoniades, C. (2019). The role of adipose tissue in cardiovascular health and disease. 0–99.

2. Chait, A., and Hartigh, L.J. den (2020). Adipose Tissue Distribution, Inflammation and Its Metabolic Consequences, Including Diabetes and Cardiovascular Disease. Frontiers in Cardiovascular Medicine 7, 22–22. 10.3389/fcvm.2020.00022.

3. Kawai, T., Autieri, M.V., and Scalia, R. (2021). Adipose tissue inflammation and metabolic dysfunction in obesity. American Journal of Physiology - Cell Physiology 320, C375–C391. 10.1152/ajpcell.00379.2020/asset/images/large/aj-acel210014f002.jpeg.

4. Neeland, I.J., Turer, A.T., Ayers, C.R., Berry, J.D., Rohatgi, A., Das, S.R., Khera, A., Vega, G.L., McGuire, D.K., Grundy, S.M., et al. (2015). Body Fat Distribution and Incident Cardiovascular Disease in Obese Adults. Journal of the American College of Cardiology 65, 2150–2151. 10.1016/j.jacc.2015.01.061.

5. Becher, T., Palanisamy, S., Kramer, D.J., Eljalby, M., Marx, S.J., Wibmer, A.G., Butler, S.D., Jiang, C.S., Vaughan, R., Schöder, H., et al. (2021). Brown adipose tissue is associated with cardiometabolic health. Nature Medicine 2021 27:1 27, 58–65. 10.1038/s41591-020-1126-7.

6. Machado, S.A., Pasquarelli-do-Nascimento, G., Silva, D.S.S. da, Farias, G.R., Santos, I. de O., Baptista, L.B., and Magalhães, K.G. (2022). Browning of the white adipose tissue regulation: new insights into nutritional and metabolic relevance in health and diseases. Nutrition & Metabolism 2022 19:1 19, 1–27. 10.1186/s12986-022-00694-0.

7. Vijay, J., Gauthier, M.F., Biswell, R.L., Louiselle, D.A., Johnston, J.J., Cheung, W.A., Belden, B., Pramatarova, A., Biertho, L., Gibson, M., et al. (2020). Single-cell analysis of human adipose tissue identifies depot- and disease-specific cell types. Nature Metabolism 2, 97–109. 10.1038/s42255-019-0152-6.

8. Rajbhandari, P., Arneson, D., Hart, S.K., Ahn, I.S., Diamante, G., Santos, L.C., Zaghari, N., Feng, A.C., Thomas, B.J., Vergnes, L., et al. (2019). Single cell analysis reveals immune cell-adipocyte crosstalk regulating the transcription of thermogenic adipocytes. eLife 8. 10.7554/elife.49501.

9. Bäckdahl, J., Franzén, L., Massier, L., Li, Q., Jalkanen, J., Gao, H., Andersson, A., Bhalla, N., Thorell, A., Rydén, M., et al. (2021). Spatial mapping reveals human adipocyte subpopulations with distinct sensitivities to insulin. Cell Metabolism 33, 1869–1882.e6. 10.1016/j.cmet.2021.07.018.

10. Karlina, R., Lutter, D., Miok, V., Fischer, D., Altun, I., Schöttl, T., Schorpp, K., Israel, A., Cero, C., Johnson, J.W., et al. (2021). Identification and characterization of distinct brown adipocyte subtypes in C57BL/6J mice. Life Science Alliance 4. 10.26508/lsa.202000924.

11. Shamsi, F., Piper, M., Ho, L.-L., Huang, T.L., Gupta, A., Streets, A., Lynes, M.D., and Tseng, Y.-H. (2021). Vascular smooth muscle-derived Trpv1+ progenitors are a source of cold-induced thermogenic adipocytes. Nature Metabolism 2021 3:4 3, 485–495. 10.1038/s42255-021-00373-z.

12. Sun, W., Dong, H., Balaz, M., Slyper, M., Drokhlyansky, E., Colleluori, G., Giordano, A., Kovanicova, Z., Stefanicka, P., Balazova, L., et al. (2020). snRNA-seq reveals a subpopulation of adipocytes that regulates thermogenesis. Nature 2020 587:7832 587, 98–102. 10.1038/s41586-020-2856-x.

13. Whytock, K.L., Sun, Y., Divoux, A., Yu, G., Smith, S.R., Walsh, M.J., and Sparks, L.M. (2022). Single cell full-length transcriptome of human subcutaneous adipose tissue reveals unique and heterogeneous cell populations. iScience 25, 104772–104772. 10.1016/j.isci.2022.104772.

14. Hill, D.A., Lim, H.W., Kim, Y.H., Ho, W.Y., Foong, Y.H., Nelson, V.L., Nguyen, H.C.B., Chegireddy, K., Kim, J., Habertheuer, A., et al. (2018). Distinct macrophage populations direct inflammatory versus physiological changes in adipose tissue. Proceedings of the National Academy of Sciences of the United States of America 115, E5096– E5105. 10.1073/pnas.1802611115.

15. Cho, D.S., Lee, B., and Doles, J.D. (2019). Refining the adipose progenitor cell landscape in healthy and obese visceral adipose tissue using single-cell gene expression profiling. Life Science Alliance 2, e201900561–e201900561. 10.26508/lsa.201900561.

16. Jaitin, D.A., Adlung, L., Thaiss, C.A., Weiner, A., Li, B., Descamps, H., Lundgren, P., Bleriot, C., Liu, Z., Deczkowska, A., et al. (2019). Lipid-Associated Macrophages Control Metabolic Homeostasis in a Trem2-Dependent Manner. Cell 178, 686–698.e14. 10.1016/j.cell.2019.05.054.

17. Harasymowicz, N.S., Rashidi, N., Savadipour, A., Wu, C.-L., Tang, R., Bramley, J., Buchser, W., and Guilak, F. (2021). Single-cell RNA sequencing reveals the induction of novel myeloid and myeloid-associated cell populations in visceral fat with long-term obesity. The FASEB Journal 35, e21417–e21417. 10.1096/fj.202001970r.

18. Hildreth, A.D., Ma, F., Wong, Y.Y., Sun, R., Pellegrini, M., and O’Sullivan, T.E. (2021). Single-cell sequencing of human white adipose tissue identifies new cell states in health and obesity. Nature Immunology, 1–15. 10.1038/s41590-021-00922-4.

19. Sárvári, A.K., Hauwaert, E.L.V., Markussen, L.K., Gammelmark, E., Marcher, A.B., Ebbesen, M.F., Nielsen, R., Brewer, J.R., Madsen, J.G.S., and Mandrup, S. (2021). Plasticity of Epididymal Adipose Tissue in Response to Diet-Induced Obesity at Single-Nucleus Resolution. Cell Metabolism 33, 437–453.e5. 10.1016/j.cmet.2020.12.004.

20. Cottam, M.A., Caslin, H.L., Winn, N.C., and Hasty, A.H. (2022). Multiomics reveals persistence of obesity-associated immune cell phenotypes in adipose tissue during weight loss and weight regain in mice. Nature Communications 2022 13:1 13, 1–16. 10.1038/s41467-022-30646-4.

21. Emont, M.P., Jacobs, C., Essene, A.L., Pant, D., Tenen, D., Colleluori, G., Vincenzo, A.D., Jørgensen, A.M., Dashti, H., Stefek, A., et al. (2022). A single-cell atlas of human and mouse white adipose tissue. Nature 603, 926– 933. 10.1038/s41586-022-04518-2.

22. McLellan, M.A., Skelly, D.A., Dona, M.S.I., Squiers, G.T., Farrugia, G.E., Gaynor, T.L., Cohen, C.D., Pandey, R., Diep, H., Vinh, A., et al. (2020). High-Resolution Transcriptomic Profiling of the Heart During Chronic Stress Reveals Cellular Drivers of Cardiac Fibrosis and Hypertrophy. Circulation 142, 1448–1463. 10.1161/circulationaha.119.045115.

23. Larsen, T.J., Jespersen, N.Z., and Scheele, C. (2018). Brown Adipose Tissue. Handb. Exp. Pharmacol. 251, 73–84. 10.1007/164_2018_142.

24. Karnam, P., Standaert, M.L., Galloway, L., and Farese, R.V. (1997). Activation and Translocation of Rho (and ADP Ribosylation Factor) by Insulin in Rat Adipocytes. Journal of Biological Chemistry 272, 6136–6140. 10.1074/jbc.272.10.6136.

25. Vergnes, L., Chin, R., Young, S.G., and Reue, K. (2011). Heart-type Fatty Acid-binding Protein Is Essential for Efficient Brown Adipose Tissue Fatty Acid Oxidation and Cold Tolerance. The Journal of Biological Chemistry 286, 380–380. 10.1074/jbc.m110.184754.

26. Cui, X., Jing, J., Wu, R., Cao, Q., Li, F., Li, K., Wang, S., Yu, L., Schwartz, G., Shi, H., et al. (2021). Adipose tissue-derived neurotrophic factor 3 regulates sympathetic innervation and thermogenesis in adipose tissue. Nature Communications 2021 12:1 12, 1–18. 10.1038/s41467-021-25766-2.

27. Subramani, M., and Yun, J.W. (2021). Loss of lymphocyte cytosolic protein 1 (LCP1) induces browning in 3T3-L1 adipocytes via β3-AR and the ERK-independent signaling pathway. The International Journal of Biochemistry & Cell Biology 138, 106053–106053. 10.1016/j.biocel.2021.106053.

28. Leonardsson, G., Steel, J.H., Christian, M., Pocock, V., Milligan, S., Bell, J., So, P.W., Medina-Gomez, G., Vidal-Puig, A., White, R., et al. (2004). Nuclear receptor corepressor RIP140 regulates fat accumulation. Proceedings of the National Academy of Sciences of the United States of America 101, 8437–8442. 10.1073/pnas.0401013101/suppl_file/01013fig7.pdf.

29. Kiefer, F.W., Vernochet, C., O’Brien, P., Spoerl, S., Brown, J.D., Nallamshetty, S., Zeyda, M., Stulnig, T.M., Cohen, D.E., Kahn, C.R., et al. (2012). Retinaldehyde dehydrogenase 1 regulates a thermogenic program in white adipose tissue. Nature medicine 18, 918–918. 10.1038/nm.2757.

30. Turpin, S.M., Nicholls, H.T., Willmes, D.M., Mourier, A., Brodesser, S., Wunderlich, C.M., Mauer, J., Xu, E., Hammerschmidt, P., Brönneke, H.S., et al. (2014). Obesity-induced CerS6-dependent C16:0 ceramide production promotes weight gain and glucose intolerance. Cell Metabolism 20, 678–686. 10.1016/j.cmet.2014.08.002.

31. Choi, K.M., Ko, C.Y., An, S.M., Cho, S.H., Rowland, D.J., Kim, J.H., Fasoli, A., Chaudhari, A.J., Bers, D.M., and Yoon, J.C. (2023). Regulation of beige adipocyte thermogenesis by the cold-repressed ER protein NNAT. Molecular Metabolism 69, 101679–101679. 10.1016/j.molmet.2023.101679.

32. Ross, S.E., Hemati, N., Longo, K.A., Bennett, C.N., Lucas, P.C., Erickson, R.L., and MacDougald, O.A. (2000). Inhibition of Adipogenesis by Wnt Signaling. Science 289, 950–950.

33. Bennett, C.N., Ross, S.E., Longo, K.A., Bajnok, L., Hemati, N., Johnson, K.W., Harrison, S.D., and MacDougald, O.A. (2002). Regulation of Wnt signaling during adipogenesis. Journal of Biological Chemistry 277, 30998–31004. 10.1074/jbc.m204527200.

34. Zamani, N., and Brown, C.W. (2011). Emerging Roles for the Transforming Growth Factor-β Superfamily in Regulating Adiposity and Energy Expenditure. Endocr. Rev. 32, 387–403. 10.1210/er.2010-0018.

35. Villanueva, C.J., Vergnes, L., Wang, J., Drew, B.G., Hong, C., Tu, Y., Hu, Y., Peng, X., Xu, F., Saez, E., et al. (2013). Adipose Subtype-Selective Recruitment of TLE3 or Prdm16 by PPARγ Specifies Lipid Storage versus Thermogenic Gene Programs. Cell Metab. 17, 423–435. 10.1016/j.cmet.2013.01.016.

36. Aibar, S., González-Blas, C.B., Moerman, T., Huynh-Thu, V.A., Imrichova, H., Hulselmans, G., Rambow, F., Marine, J.C., Geurts, P., Aerts, J., et al. (2017). SCENIC: Single-cell regulatory network inference and clustering. Nature methods 14, 1083–1083. 10.1038/nmeth.4463.

37. Randi, A.M., Sperone, A., Dryden, N.H., and Birdsey, G.M. (2009). Regulation of angiogenesis by ETS transcription factors. Biochemical Society Transactions 37, 1248–1253. 10.1042/bst0371248.

38. Deleuze, V., El-Hajj, R., Chalhoub, E., Dohet, C., Pinet, V., Couttet, P., and Mathieu, D. (2012). Angiopoietin-2 is a direct transcriptional target of TAL1, LYL1 and LMO2 in endothelial cells. PloS one 7. 10.1371/journal.pone.0040484.

39. Rehli, M., Sulzbacher, S., Pape, S., Ravasi, T., Wells, C.A., Heinz, S., Söllner, L., Chartouni, C.E., Krause, S.W., Steingrimsson, E., et al. (2005). Transcription factor Tfec contributes to the IL-4-inducible expression of a small group of genes in mouse macrophages including the granulocyte colony-stimulating factor receptor. Journal of immunology (Baltimore, Md. : 1950) 174, 7111–7122. 10.4049/jimmunol.174.11.7111.

40. Krausgruber, T., Blazek, K., Smallie, T., Alzabin, S., Lockstone, H., Sahgal, N., Hussell, T., Feldmann, M., and Udalova, I.A. (2011). IRF5 promotes inflammatory macrophage polarization and TH1-TH17 responses. Nature Immunology 2011 12:3 12, 231–238. 10.1038/ni.1990.

41. Alexanian, M., Przytycki, P.F., Micheletti, R., Padmanabhan, A., Ye, L., Travers, J.G., Gonzalez-Teran, B., Silva, A.C., Duan, Q., Ranade, S.S., et al. (2021). A transcriptional switch governs fibroblast activation in heart disease. Nature 2021 595:7867 595, 438–443. 10.1038/s41586-021-03674-1.

42. Khalil, H., Kanisicak, O., Prasad, V., Correll, R.N., Fu, X., Schips, T., Vagnozzi, R.J., Liu, R., Huynh, T., Lee, S.J., et al. (2017). Fibroblast-specific TGF-β-Smad2/3 signaling underlies cardiac fibrosis. The Journal of clinical investigation 127, 3770–3783. 10.1172/jci94753.

43. Cinti, S., Mitchell, G., Barbatelli, G., Murano, I., Ceresi, E., Faloia, E., Wang, S., Fortier, M., Greenberg, A.S., Obin, M.S., et al. (2005). Adipocyte death defines macrophage localization and function in adipose tissue of obese mice and humans. Journal of Lipid Research 46, 2347–2355. 10.1194/jlr.m500294-jlr200.

44. Amano, S.U., Cohen, J.L., Vangala, P., Tencerova, M., Nicoloro, S.M., Yawe, J.C., Shen, Y., Czech, M.P., and Aouadi, M. (2014). Local proliferation of macrophages contributes to obesity-associated adipose tissue inflammation. Cell metabolism 19, 162–162. 10.1016/j.cmet.2013.11.017.

45. Zheng, C., Yang, Q., Cao, J., Xie, N., Liu, K., Shou, P., Qian, F., Wang, Y., and Shi, Y. (2016). Local proliferation initiates macrophage accumulation in adipose tissue during obesity. Cell Death & Disease 2016 7:3 7, e2167–e2167. 10.1038/cddis.2016.54.

46. Eguchi, J., Kong, X., Tenta, M., Wang, X., Kang, S., and Rosen, E.D. (2013). Interferon Regulatory Factor 4 Regulates Obesity-Induced Inflammation Through Regulation of Adipose Tissue Macrophage Polarization. Diabetes 62, 3394–3403. 10.2337/db12-1327.

47. Sardiello, M., Palmieri, M., Ronza, A.D., Medina, D.L., Valenza, M., Gennarino, V.A., Malta, C.D., Donaudy, F., Embrione, V., Polishchuk, R.S., et al. (2009). A gene network regulating lysosomal biogenesis and function. Science 325, 473–477. 10.1126/science.1174447/suppl_file/sardiello-som.pdf.

48. He, Y.S., Yang, X.K., Hu, Y.Q., Xiang, K., and Pan, H.F. (2021). Emerging role of Fli1 in autoimmune diseases. International Immunopharmacology 90, 107127–107127. 10.1016/j.intimp.2020.107127.

49. McGinnis, C.S., Patterson, D.M., Winkler, J., Conrad, D.N., Hein, M.Y., Srivastava, V., Hu, J.L., Murrow, L.M., Weissman, J.S., Werb, Z., et al. (2019). MULTI-seq: sample multiplexing for single-cell RNA sequencing using lipid-tagged indices. Nature Methods 2019 16:7 16, 619–626. 10.1038/s41592-019-0433-8.

50. Yoshida, T., Nishioka, H., Yoshioka, K., and Kondo, M. (1987). Reduced norepinephrine turnover in interscapular brown adipose tissue of obese rats after ovariectomy. Metabolism 36, 1–6. 10.1016/0026-0495(87)90054-0.

51. Yoshioka, K., Yoshida, T., Wakabayashi, Y., Nishioka, H., and Kondo, M. (1988). Reduced Brown Adipose Tissue Thermogenesis of Obese Rats After Ovariectomy. Endocrinologia Japonica 35, 537–543. 10.1507/endocrj1954.35.537.

52. Quevedo, S., Roca, P., Picó, C., and Palou, A. (1998). Sex-associated differences in cold-induced UCP1 synthesis in rodent brown adipose tissue. Pflügers Archiv 1998 436:5 436, 689–695. 10.1007/s004240050690.

53. Justo, R., Frontera, M., Pujol, E., Rodríguez-Cuenca, S., Lladó, I., García-Palmer, F.J., Roca, P., and Gianotti, M. (2005). Gender-related differences in morphology and thermogenic capacity of brown adipose tissue mitochondrial subpopulations. Life Sciences 76, 1147–1158. 10.1016/j.lfs.2004.08.019.

54. Kim, S.N., Jung, Y.S., Kwon, H.J., Seong, J.K., Granneman, J.G., and Lee, Y.H. (2016). Sex differences in sympathetic innervation and browning of white adipose tissue of mice. Biology of Sex Differences 7, 1–13. 10.1186/s13293-016-0121-7.

55. Rudnicki, M., Abdifarkosh, G., Rezvan, O., Nwadozi, E., Roudier, E., and Haas, T.L. (2018). Female mice have higher angiogenesis in perigonadal adipose tissue than males in response to high-fat diet. Frontiers in Physiology 9, 1452–1452. 10.3389/fphys.2018.01452.

56. Bagchi, D.P., Nishii, A., Li, Z., DelProposto, J.B., Corsa, C.A., Mori, H., Hardij, J., Learman, B.S., Lumeng, C.N., and MacDougald, O.A. (2020). Wnt/β-catenin signaling regulates adipose tissue lipogenesis and adipocyte-specific loss is rigorously defended by neighboring stromal-vascular cells. Molecular metabolism 42. 10.1016/j.molmet.2020.101078.

57. Palani, N.P., Horvath, C., Timshel, P.N., Folkertsma, P., Grønning, A.G.B., Henriksen, T.I., Peijs, L., Jensen, V.H., Sun, W., Jespersen, N.Z., et al. (2023). Adipogenic and SWAT cells separate from a common progenitor in human brown and white adipose depots. Nature Metabolism 2023 5:6 5, 996–1013. 10.1038/s42255-023-00820-z.

58. Haase, J., Weyer, U., Immig, K., Klöting, N., Blüher, M., Eilers, J., Bechmann, I., and Gericke, M. (2014). Local proliferation of macrophages in adipose tissue during obesity-induced inflammation. Diabetologia 57, 562–571. 10.1007/s00125-013-3139-y/figures/7.

59. Zamarron, B.F., Mergian, T.A., Cho, K.W., Martinez-Santibanez, G., Luan, D., Singer, K., DelProposto, J.L., Geletka, L.M., Muir, L.A., and Lumeng, C.N. (2017). Macrophage Proliferation Sustains Adipose Tissue Inflammation in Formerly Obese Mice. Diabetes 66, 392–406. 10.2337/db16-0500.

60. Palmieri, M., Impey, S., Kang, H., Ronza, A. di, Pelz, C., Sardiello, M., and Ballabio, A. (2011). Characterization of the CLEAR network reveals an integrated control of cellular clearance pathways. Human Molecular Genetics 20, 3852–3866. 10.1093/hmg/ddr306.

61. Mizunoe, Y., Kobayashi, M., Hoshino, S., Tagawa, R., Itagawa, R., Hoshino, A., Okita, N., Sudo, Y., Nakagawa, Y., Shimano, H., et al. (2020). Cathepsin B overexpression induces degradation of perilipin 1 to cause lipid metabolism dysfunction in adipocytes. Scientific Reports 2020 10:1 10, 1–12. 10.1038/s41598-020-57428-6.

62. Mizunoe, Y., Sudo, Y., Okita, N., Hiraoka, H., Mikami, K., Narahara, T., Negishi, A., Yoshida, M., Higashibata, R., Watanabe, S., et al. (2017). Involvement of lysosomal dysfunction in autophagosome accumulation and early pathologies in adipose tissue of obese mice. Autophagy 13, 642–653. 10.1080/15548627.2016.1274850.

63. Haka, A.S., Barbosa-Lorenzi, V.C., Lee, H.J., Falcone, D.J., Hudis, C.A., Dannenberg, A.J., and Maxfield, F.R. (2016). Exocytosis of macrophage lysosomes leads to digestion of apoptotic adipocytes and foam cell formation[S]. J. Lipid Res. 57, 980–992. 10.1194/jlr.m064089.

64. Rodríguez, E., Monjo, M., Rodríguez-Cuenca, S., Pujol, E., Amengual, B., Roca, P., and Palou, A. (2014). Sexual dimorphism in the adrenergic control of rat brown adipose tissue response to overfeeding. Pflügers Archiv 2001 442:3 442, 396–403. 10.1007/s004240100556.

65. Rodríguez-Cuenca, S., Pujol, E., Justo, R., Frontera, M., Oliver, J., Gianotti, M., and Roca, P. (2002). Sex-dependent thermogenesis, differences in mitochondrial morphology and function, and adrenergic response in brown adipose tissue. Journal of Biological Chemistry 277, 42958–42963. 10.1074/jbc.m207229200.

66. Rodríguez, A.M., Quevedo-Coli, S., Roca, P., and Palou, A. (2001). Sex-dependent dietary obesity, induction of UCPs, and leptin expression in rat adipose tissues. Obesity Research 9, 579–588. 10.1038/oby.2001.75.

67. Cinti, S., Cancello, R., Zingaretti, M.C., Ceresi, E., Matteis, R.D., Giordano, A., Himms-Hagen, J., and Ricquier, D. (2002). CL316,243 and cold stress induce heterogeneous expression of UCP1 mRNA and protein in rodent brown adipocytes. Journal of Histochemistry and Cytochemistry 50, 21–31. 10.1177/002215540205000103/asset/images/large/10.1177_002215540205000103-fig2.jpeg.

68. Song, A., Dai, W., Jang, M.J., Medrano, L., Li, Z., Zhao, H., Shao, M., Tan, J., Li, A., Ning, T., et al. (2020). Low- And high-thermogenic brown adipocyte subpopulations coexist in murine adipose tissue. Journal of Clinical Investigation 130, 247–257. 10.1172/jci129167.

69. Shimizu, I., Aprahamian, T., Kikuchi, R., Shimizu, A., Papanicolaou, K.N., MacLauchlan, S., Maruyama, S., and Walsh, K. (2014). Vascular rarefaction mediates whitening of brown fat in obesity. The Journal of Clinical Investigation 124, 2099–2099. 10.1172/jci71643.

70. Fatima, L.A., Campello, R.S., Santos, R.D.S., Freitas, H.S., Frank, A.P., Machado, U.F., and Clegg, D.J. (2017). Estrogen receptor 1 (ESR1) regulates VEGFA in adipose tissue. Scientific Reports 2017 7:1 7, 1–14. 10.1038/s41598-017-16686-7.

71. Suriano, F., Vieira-Silva, S., Falony, G., Roumain, M., Paquot, A., Pelicaen, R., Régnier, M., Delzenne, N.M., Raes, J., Muccioli, G.G., et al. (2021). Novel insights into the genetically obese (ob/ob) and diabetic (db/db) mice: two sides of the same coin. Microbiome 9, 1–20. 10.1186/s40168-021-01097-8/figures/7.

72. Jong, J.M.A. de, Sun, W., Pires, N.D., Frontini, A., Balaz, M., Jespersen, N.Z., Feizi, A., Petrovic, K., Fischer, A.W., Bokhari, M.H., et al. (2019). Human brown adipose tissue is phenocopied by classical brown adipose tissue in physiologically humanized mice. Nature Metabolism 1, 830–843. 10.1038/s42255-019-0101-4.

73. Huang, Z., Zhang, Z., Moazzami, Z., Zhu, W., Shen, S., and Ruan, H.-B. (2022). Brown adipose tissue involution associated with progressive restriction in progenitor competence. CellReports 39, 110575–110575. 10.1016/j.celrep.2022.110575.

74. Coleman, D.L. (1982). Diabetes-Obesity Syndromes in Mice. Diabetes 31, 1–6. 10.2337/diab.31.1.s1.

75. Coleman, D.L. (1978). Obese and diabetes: Two mutant genes causing diabetes-obesity syndromes in mice. Diabetologia 14, 141–148. 10.1007/bf00429772.

76. Chen, H., Charlat, O., Tartaglia, L.A., Woolf, E.A., Weng, X., Ellis, S.J., Lakey, N.D., Culpepper, J., More, K.J., Breitbart, R.E., et al. (1996). Evidence That the Diabetes Gene Encodes the Leptin Receptor: Identification of a Mutation in the Leptin Receptor Gene in db/db Mice. Cell 84, 491–495. 10.1016/s0092-8674(00)81294-5.

77. Pinto, A.R., Chandran, A., Rosenthal, N.A., and Godwin, J.W. (2013). Isolation and analysis of single cells from the mouse heart. Journal of Immunological Methods 393, 74–80. 10.1016/j.jim.2013.03.012.

78. Zhu, C., Yu, M., Huang, H., Juric, I., Abnousi, A., Hu, R., Lucero, J., Behrens, M.M., Hu, M., and Ren, B. (2019). An ultra high-throughput method for single-cell joint analysis of open chromatin and transcriptome. Nat. Struct. Mol. Biol. 26, 1063–1070. 10.1038/s41594-019-0323-x.

79. Hafemeister, C., and Satija, R. (2019). Normalization and variance stabilization of single-cell RNA-seq data using regularized negative binomial regression. Genome Biology 20, 1–15. 10.1186/s13059-019-1874-1/figures/6.

80. McGinnis, C.S., Murrow, L.M., and Gartner, Z.J. (2019). DoubletFinder: Doublet Detection in Single-Cell RNA Sequencing Data Using Artificial Nearest Neighbors. Cell Systems 8, 329–337.e4. 10.1016/j.cels.2019.03.003.

81. Finak, G., McDavid, A., Yajima, M., Deng, J., Gersuk, V., Shalek, A.K., Slichter, C.K., Miller, H.W., McElrath, M.J., Prlic, M., et al. (2015). MAST: A flexible statistical framework for assessing transcriptional changes and characterizing heterogeneity in single-cell RNA sequencing data. Genome Biology 16, 1–13. 10.1186/s13059-015-0844-5/figures/6.

82. Soneson, C., and Robinson, M.D. (2018). Bias, robustness and scalability in single-cell differential expression analysis. Nature Methods 2018 15:4 15, 255–261. 10.1038/nmeth.4612.

83. Yu, G., Wang, L.G., Han, Y., and He, Q.Y. (2012). ClusterProfiler: An R package for comparing biological themes among gene clusters. OMICS A Journal of Integrative Biology 16, 284–287. 10.1089/omi.2011.0118/asset/images/large/figure1.jpeg.

84. Schindelin, J., Arganda-Carreras, I., Frise, E., Kaynig, V., Longair, M., Pietzsch, T., Preibisch, S., Rueden, C., Saalfeld, S., Schmid, B., et al. (2012). Fiji: an open-source platform for biological-image analysis. Nature Methods 2012 9:7 9, 676–682. 10.1038/nmeth.2019.

